# Targeting ion channel TRPM7 promotes thymic development of Regulatory T cells by increasing IL-2-dependent STAT5 activation

**DOI:** 10.1101/330233

**Authors:** Suresh K. Mendu, Michael S. Schappe, Emily K. Moser, Julia K. Krupa, Jason S. Rogers, Eric J. Stipes, Clare A. Parker, Thomas J. Braciale, Bimal N. Desai

## Abstract

**In Brief:** Genetic deletion of *Trpm7* in T-cells or pharmacological inhibition of TRPM7 channel promotes the development of fully functional T_reg_ cells by increasing IL-2Rα and STAT5-dependent FOXP3 expression in the developing thymocytes. The study identifies the ion channel TRPM7 as a putative drug target to increase T_reg_ numbers *in vivo* and induce immunotolerance.

**HIGHLIGHTS:**

- Ion channel TRPM7 controls T_reg_ development
- The deletion of *Trpm7* in the T-cell lineage increases fully functional T_reg_ cells in the periphery
- TRPM7 negatively regulates *Foxp3* expression by restraining IL-2-dependent STAT5 activation
- Inhibition of TRPM7 channel by FTY720 promotes the development of functional T_reg_ cells

**SUMMARY:** The thymic development of regulatory T cells (T_reg_), the crucial suppressors of the effector T cells (T_eff_), is governed by the transcription factor FOXP3. Despite the clinical significance of T_reg_ cells, there is a dearth of druggable molecular targets capable of increasing T_reg_ numbers *in vivo*. We report a surprising discovery that TRPM7 restrains T_reg_ development by negatively regulating STAT5-dependent *Foxp3* expression. The deletion of *Trpm7* potentiates the thymic development of T_reg_ cells, leads to a significantly higher frequency of functional T_reg_ cells in the periphery and renders the mice highly resistant to T cell-dependent hepatitis. The deletion of *Trpm7* or the inhibition of TRPM7 channel activity by the FDA-approved prodrug FTY720, increases IL-2 sensitivity through a feed forward positive feedback loop involving high IL-2Rα expression and STAT5 activation. Enhanced IL-2 signaling increases the expression of *Foxp3* in thymocytes and promotes the development of T_reg_ cells. Thus, TRPM7 emerges as the first ion channel that can be drugged to increase T_reg_ numbers, revealing a novel pharmacological path toward the induction of immune tolerance.

## INTRODUCTION

The immune system has the ability to control the intensity, duration and scope of inflammatory processes through an elaborate array of checks and balances (Germain, 2012). Regulatory T cells (T_reg_) play a salient immunosuppressive role to balance the destructive potential of effector T cells (T_eff_) (Ramsdell and Ziegler, 2014). Although T_reg_ cells develop from the same thymocyte precursors as T_eff_ cells, the distinct T_reg_ lineage is specified through the expression and maintenance of the transcription factor FOXP3 (Hori et al., 2003). Consequently, the genetic loss of *FOXP3* in humans and mice prevents the development of T_reg_ cells, and breaks self-tolerance (Bennett et al., 2001; Brunkow et al., 2001; Fontenot et al., 2003; Khattri et al., 2003; Wildin et al., 2001). Interleukin-2 (IL-2) plays a decisive role in immune tolerance by regulating the development, maintenance and function of T_reg_ cells (Bayer et al., 2013; Fontenot et al., 2005; Malek, 2003). The key transcription factor responsible for *Foxp3* gene expression, downstream of IL-2 signaling, is STAT5 (Antov et al., 2003; Burchill et al., 2007) and accordingly, ectopic expression of a constitutively active STAT5 variant is sufficient to divert the fate of developing thymocytes toward the T_reg_ lineage (Burchill et al., 2008). The thymic development of T_reg_ cells has been theorized to occur in a two-step process (Lio and Hsieh, 2008). In the first step, T cell receptor (TCR) signaling upregulates the IL-2Rα (CD25) and other components of IL-2 signaling pathway, increasing the sensitivity of IL-2 signaling. In the second step, IL-2 mediated signals upregulate *Foxp3* in a STAT5-dependent manner to finalize the commitment to T_reg_ lineage. Identifying novel “druggable” components in this pathway may enable the pharmacological manipulation of *Foxp3* expression and T_reg_ numbers in vivo.

The National Center for Advancing Translational Sciences (NCATS) has identified a druggable genome of ∼3000 human genes encoding predominantly three key protein families that are ideal for drug development: non-olfactory GPCRs, ion channels and protein kinases. Included in the ion channel genes are the <u>T</u>ransient <u>r</u>eceptor <u>p</u>otential (TRP) channels, a 28-member superfamily of ion channels that constitute an exciting class of drug targets (Nilius and Szallasi, 2014). Cation-selective TRP channels mediate context-specific electrical signaling in all organ systems but the functions of the specific TRP channels prevalent in the immune system remain largely mysterious and thus unexploited for immunomodulation. Three members of the TRP superfamily: TRPM2, TRPM6 and TRPM7 are referred to as *chanzymes* because they possess ion <u>chan</u>nel activity as well as an additional en<u>zyme</u> activity. TRPM7, an ion channel permeable to cations such as Ca^2+^, Na^+^, Zn^2+^ and Mg^2+^, contains a carboxy-terminal serine-threonine kinase domain (Monteilh-Zoller et al., 2003; Nadler et al., 2001; Runnels et al., 2001) and is highly expressed in the T-cell lineage (Jin et al., 2008). Previously, we generated *Trpm7*^*fl/fl*^*(Lck Cre)* mice to delete *Trpm7* selectively in the T-cell lineage (Jin et al., 2008). In these mice, thymocyte development is impaired resulting in a significant accumulation of CD4-CD8-thymocytes and reduced thymic cellularity (Jin et al., 2008). The developmental block is partial, and the residual egress of mature CD4+ and CD8+ T cells populates the peripheral lymphoid organs normally. Since *Trpm7-/-* T cells are also resistant to apoptosis (Desai et al., 2012), we expected to see a lymphoproliferative phenotype and autoimmunity (Bidere et al., 2006). Surprisingly, the *Trpm7*^*fl/fl*^ *(Lck Cre)* mice [hereafter, “KO” mice], exhibit T-cell lymphopenia (Jin et al., 2008) and autoimmunity is greatly delayed and mild in its manifestation. Lung inflammation is seen in 12 month old KO mice (Desai et al., 2012). These observations suggest that deletion of TRPM7 in T-cells promotes immunosuppression through an undefined mechanism.

In the present study, we studied an autoimmune hepatitis (AIH) model, a T-cell dependent inflammatory disease that often leads to liver cirrhosis, cancer and death (Manns et al., 2015). In mice, AIH is modeled by intravenous (i.v.) injection of Concanavalin A (Con A) which results in an acute dose-dependent liver inflammation that is dependent on the activity of CD4+ T cells (Tiegs et al., 1992). Through these studies, we have made a serendipitous discovery that the deletion of *Trpm7* in the T-cell lineage greatly potentiates the thymic development of T_reg_ cells and leads to a significantly higher frequency of functional T_reg_ cells in the periphery. Deletion of *Trpm7* or pharmacological inhibition of TRPM7 using the FDA-approved drug FTY720, also known as Fingolimod, and repurposed here to block TRPM7, greatly increases IL-2 dependent STAT5 activation. This increases FOXP3 expression in developing thymocytes, steering a larger percentage toward the T_reg_ lineage. Blocking TRPM7 channel activity thus emerges as an attractive pharmacological strategy to increase T_reg_ frequency *in vivo* and induce tolerance in autoimmune diseases.

## RESULTS

### Deletion of *Trpm7* in T cells protects the mice from Concanavalin A-induced experimental lethal hepatitis

Autoimmune hepatitis, an inflammatory disease of the liver (Manns et al., 2015), is modeled in mice by injecting the mice with Con A, resulting in acute T-cell dependent liver inflammation and lethality (Tiegs et al., 1992). We administered Con A (40 mg/kg, iv) and assessed the development of hepatitis over a course of 24h; the experimental design is schematized **(Fig. 1A)**. As shown in the Kaplan-Meier survival analysis of *Trpm7*^*fl/fl*^ (WT) and *Trpm7*^*fl/fl*^*(Lck Cre)* (KO) mice **(Fig. 1B)**, 50% of the WT mice (n=10) succumbed within 24h. In striking contrast, the KO mice (n=10) were completely resistant to Con A-induced death. To assess liver damage, we measured the concentration of the liver enzyme, Alanine transaminase (ALT), in the serum, 24h after Con A injections. As shown **(Fig. 1C)**, the mean serum ALT activity in WT mice increased 9-fold (120 IU/L to 1093 IU/L). However, the liver damage in Con A-injected KO mice was modest, showing only a 3-fold increase in mean serum ALT activity (134 IU/L to 398 IU/L). Through histological analysis of liver sections, perivascular inflammation was readily evident in Con A-treated WT mice but not in Con A-treated KO mice **(Fig. 1D)**. We also measured the relative changes in the gene expression of inflammatory cytokines in homogenized liver tissue, 24h after Con A injections. In the WT mice, the mRNA levels of inflammatory cytokines increased substantially in Con A-injected mice when compared to saline-injected controls **(Fig. 1E)**. For example, the mRNA levels of *Ifng* increased 250-fold and *Tnf* mRNA levels increased 70-fold. The *Il2*, *Il1b* and *Il6* gene expression were also upregulated but to a lesser extent. Strikingly, in contrast to WT mice, the increase in inflammatory cytokine expression in Con A-injected KO mice was prominently blunted. Interestingly, the trend was reversed in the case of anti-inflammatory cytokine *Tgfb1.* In the WT mice, the Con A-injected mice showed only a 2-fold increase in *Tgfb1* mRNA levels in comparision to saline-injected mice. In contrast, the *Tgfb1* mRNA levels increased 10-fold in Con A-injected KO mice, relative to saline controls. Overall, the results presented in **Figure 1** indicate that the presence of TRPM7 in T cells is essential for the onset of acute liver inflammation induced by Con A. Since the deletion of *Trpm7* in T cells is dramatically protective, we tested the hypothesis that TRPM7 is essential for T-cell activation. Indeed, this idea has been suggested previously, albeit without the benefit of gene targeted mice (Sahni and Scharenberg, 2008).

**Figure 1.**
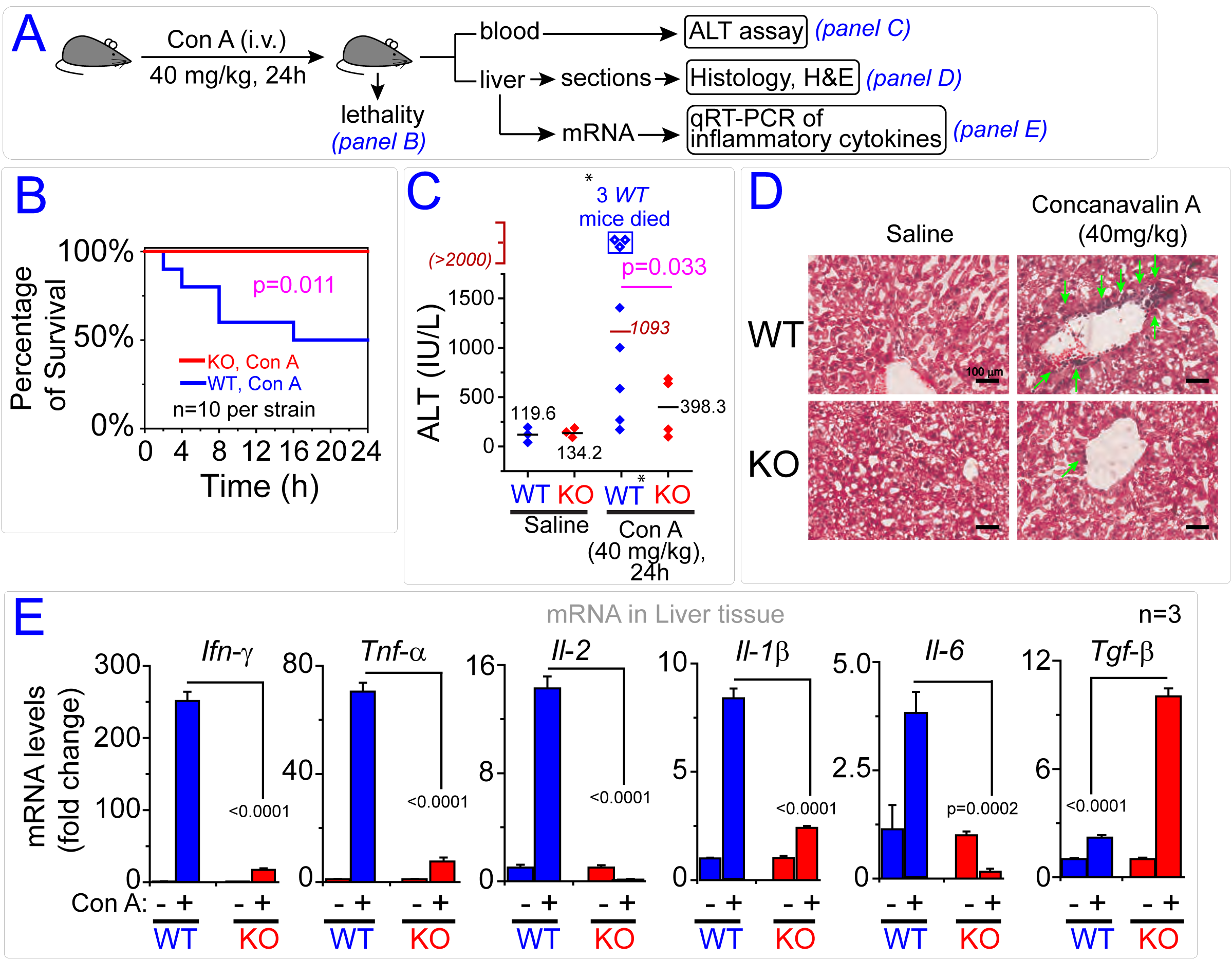
Deletion of Trpm7 in T cells protects the mice from Concanavalin A-induced experimental lethal hepatitis. **(A)** Experimental scheme. **(B)** Kaplan-Meier survival analysis of *Trpm7*^*fl/fl*^ (WT) and *Trpm7*^*fl/fl*^*(Lck Cre)* (KO) mice injected with Concanavalin A (40 mg/kg, iv) for 24h. (Statistics: Log-rank value = 6.36, p = 0.011 and n=10 mice in each condition). **(C)** Serum alanine transaminase (ALT) activity in WT and KO mice, 24h after administration of Concanavalin A or saline. For the 3 wt mice that died before 12h, the ALT levels were recorded as 2000 IU/L to calculate the mean. The data were collected from 4 independent experiments (p = 0.033, calcuated by two-tailed t-test). **(D)** Hematoxylin and Eosin (H&E) stain of liver sections from WT and KO mice injected with saline or Concanavalin A (40 mg/kg, iv). Scale bar = 100 μm. The green arrows mark the perivascular infiltration by immune cells. **(E)** Gene expression analysis (qPCR) of indicated cytokines in WT and KO liver tissue, 24h after saline or Concanavalin A administration. The data are representative of three independent expeirments. The p values were calcuated by two-tailed t-test and shown in each panel.

### *Trpm7-/-* T_eff_ cells are not deficient in T-cell activation and proliferation

T-cell activation is initiated after the stimulation of TCR complex in conjunction with a costimulatory signal (Smith-Garvin et al., 2009). To evaluate T-cell activation, we stimulated freshly isolated CD4+CD25-T cells, referred to as effector T cells (T_eff_) with either Con A or a mixture of anti-CD3 and anti-CD28 antibodies. The cells were analyzed for the activation markers by flow cytometry and for cytokine secretion by ELISA **(Fig. 2A)**. After 48h of stimulation, both WT and *Trpm7-/-* (KO) T_eff_ cells showed a robust increase in the cell surface levels of CD69 **(Fig. 2B)**. The upregulation of cell surface CD25 levels were considerably higher in the KO mice **(Fig. 2C)**, and this was a reflection of increased transcriptional output of CD25 mRNA *(discussed later)*. T-cell activation results in increased cell size which is conveniently measured by the forward scatter parameter (FSC) in flow cytometry. As shown **(Fig. 2D)**, both WT and KO T_eff_ cells increase in size comparably. After 72h of activation, we measured T-cell proliferation using the CFSE labeling method, which measures the progressive dilution of CFSE during cell divisions. Both WT and KO T_eff_ cells proliferated comparably in response to stimulation by the anti-CD3 and anti-CD28 antibody cocktail **(Fig. 2E)**. We then considered the possibility that the T cells migrate abnormally in the absence of TRPM7. We compared transwell migration of WT and KO T cells in response to chemotactic factors produced by activated macrophages. As shown **(Fig. S2A)** the KO T cells are not deficient in migration through the membrane (pore size: 5.0 μm). Since a major function of activated CD4+ T cells is the secretion of cytokines that orchestrate inflammation, we measured the concentrations of various inflammatory cytokines in the 72h culture supernatants of activated T cells. In the case of Con A stimulation, the overall secretion of cytokines was comparable between WT and KO T cells; the KO T cells secreted slightly higher levels of IL-4 but lower levels of IL-6 **(Fig. 2F)**. When the activation was carried out using a mixture of anti-CD3 and anti-CD28 antibodies, the KO T cells consistently secreted higher concentrations of IL-2, IL-4, IL-5 and IL-6 relative to WT T cells **(Fig. 2G)**. Importantly, the KO T cells also secreted higher levels of anti-inflammatory cytokine IL-10 and we confirmed that this was due to the increased transciption of *Il-10* mRNA **(Fig S2B)**. In their totality, the results in **Figure 2** and **Figure S2**, disproved our original hypothesis that the protection against Con A-induced lethal hepatitis was due to a defect in the activation or proliferation of *Trpm7*-/-T_eff_ cells. In search for explanatory clues, we characterized the composition of hematopoietic infiltration in the livers of Con A-injected mice.

**Figure 2.**
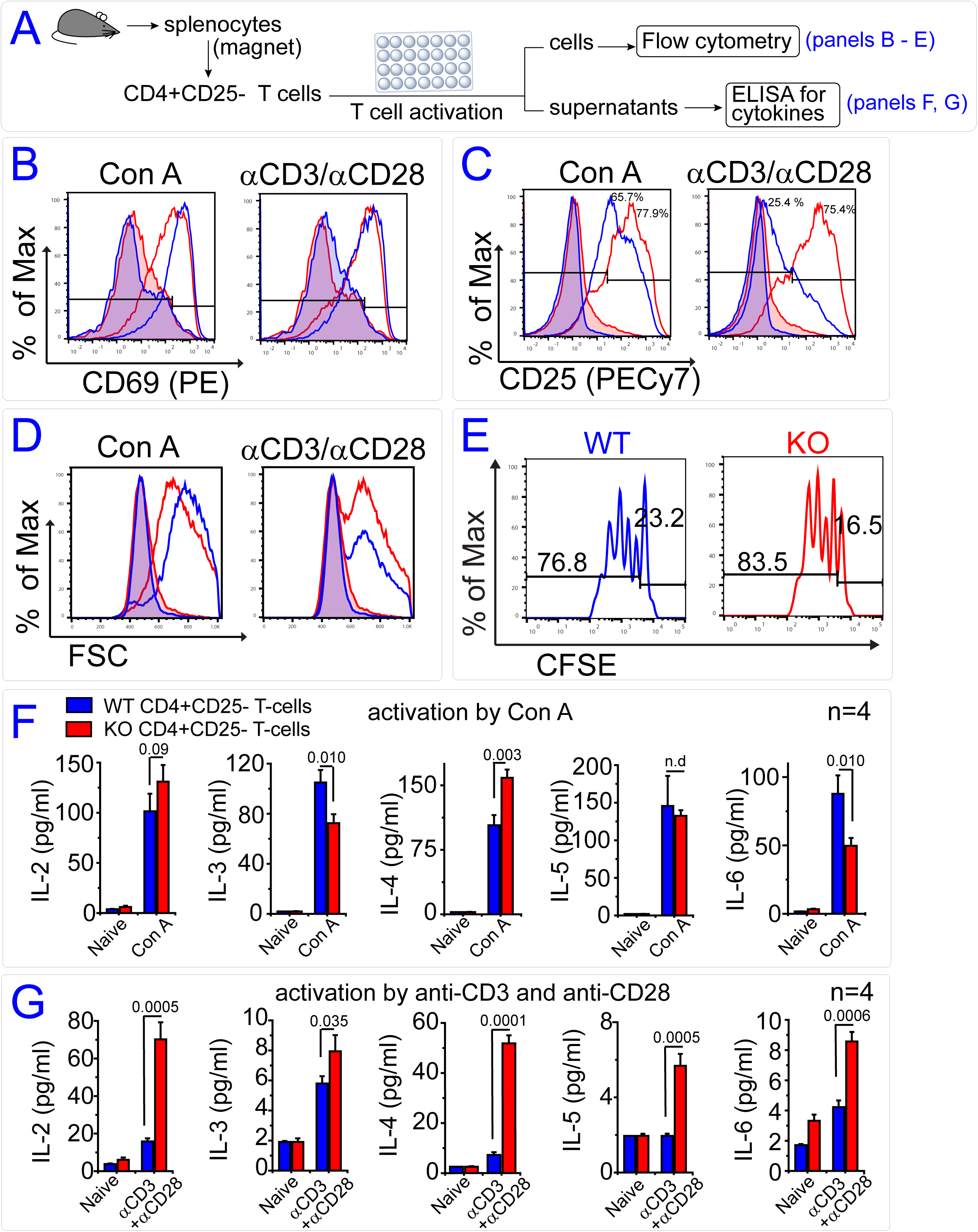
*Trpm7-/-* T_eff_ cells activate and function normally upon TCR stimulation. **(A)** Experimental scheme. **(B)** Overlaid flow cytometry histograms showing the induction of activation marker CD69 on CD4+CD25-T cells isolated from *Trpm7*^*fl/fl*^ (WT) and *Trpm7*^*fl/fl*^*(LckCre)* (KO) mice after *in vitro* activation of T cells by Concanavalin A (Con A, 5 μg/ml, 48h; *left panel*) or by a mixture of anti-CD3 and anti-CD28 antibodies (48h; *right panel*). The filled histograms are of naive cells shown in blue (WT) and red (KO); the overlap results in purple. Histograms, after activation, are shown as unfilled blue (WT) and unfilled red (KO). The flow cytometry histograms are representative of three independent experiments. **(C)** Induction of activation marker CD25 on WT and KO CD4+CD25-T cells after activation *(format and color scheme as in panel A)*. The flow cytomety histograms are representative of three independent experiments. **(D)** Increase in the size of WT and KO CD4+CD25-T cells, as measured by forward scatter (FSC) parameter after T-cell activation. The flow cytomety histograms are representative of three individual experiments. **(E)** Cell division of CFSE-labeled WT and KO CD4+CD25-T cells as measured by dilution of CFSE, 72h after activation by anti-CD3 and anti-CD28 antibodies. The percentage of dividing cells (lower gate) and undivided cells (upper gate) are quantified. The CFSE dilution histograms are representative of four independent experiments. **(F)** Quantification of secreted cytokines as detected in the supernatants of activated CD4+CD25-T cells (Con A, 5 μg/ml, 72h). The p values were calcuated by two-tailed t-test and denoted on bar graph. **(G)** Quantification of secreted cytokines as detected in the supernatants of activated CD4+CD25-T cells (anti-CD3 + anti-CD28 antibodies, 5 μg/ml, 72h). The p values were calcuated by two-tailed t-test and denoted on each bar graph.

### Increased infiltration of regulatory T cells in the livers of Con A-treated Trpm7^fl^ (*Lck Cre*) mice

The liver constitutes an unique immunological environment, housing a large number and diversity of hematopoietic cells (Heymann and Tacke, 2016). To quantify the composition of these cells by flow cytometry, we analyzed single cell suspensions from freshly excised livers, and also evaluated liver sections using immunohistochemistry and Scanning Electron Microscopy (SEM); the overall experimental scheme is illustrated **(Fig. 3A)**. Based on the cell surface staining of CD45, a marker of hematopoietic cells, the livers of saline-injected WT and KO mice contained a comparable proportion of hematopoietic cells (∼40%). A robust increase in CD45+ hematopoietic cells was seen in both WT and KO mice 24h after Con A-injections (∼80%) **(Fig. 3B)**; these data are quantified as a scatter plot (**Fig. 3C)**. This analysis demonstrates that there is no significant difference in WT and KO mice in terms of overall infiltration of hematopoietic cells following Con A injections. Since CD4+ T cells have been shown to be the key mediators of Con A-induced lethality (Tiegs et al., 1992), we assessed the percentage of CD4+ T cells in the hematopoietic cell population. As shown **(Fig. 3D)**, the liver-resident CD4+ T cells are readily detectable in saline and Con A injected mice. The scatter plot **(Fig. 3E)** shows that the percentage of CD4+ T cells in the liver-resident immune cells is modestly lower in saline-injected KO mice when compared to the WT controls. Although Con A treatment increases T cell infiltration in the liver, the percentage of CD4+ T cells is reduced due to the increased infiltration of other cell types. The gating strategy used to quantify the frequency of FOXP3+ T_reg_ cells is shown **(Fig. S3)**. The percentage of T cells that are FOXP3+ (T_reg_) is similar in the livers of saline-injected WT and KO mice **(Figures 3F** and **3G)**. Surprisingly, in Con A-injected KO mice, the percentage of FOXP3+ cells is significantly increased relative to Con A-injected WT mice (n=4; WT 21.5±4.1% and KO 39.8±3.6%; p=0.03). This result indicates that in the livers of KO mice, newly recruited T cells contain a significantly higher percentage of FOXP3+ T_reg_ cells. Since FOXP3+ T_reg_ cells are highly immunosupressive, these data strongly suggest that the increased frequency of T_reg_ cells in the KO mice protects these mice from Con A-induced liver inflammation. The lack of liver inflammation in KO mice is borne out further by the evidence that the liver sections of Con A-injected WT mice, but not KO mice, show prominent perivascular patches of CD4+ T cells in immunofluorescent confocal microscopy images **(Fig. 3H)**. Analysis of the liver vasculature by SEM revealed a large presence of endothelium-adhesive immune cells in the livers of Con A-injected WT mice, but this was clearly absent in the Con A-injected KO mice **(Fig. 3I)**. Taken together, the results presented in **Figures 3** and **S3** led us to hypothesize that the deletion of *Trpm7* in the T-cell lineage results in a significantly increased frequency of T_reg_ cells in the periphery. The decrease in T_eff_:T_reg_ ratio may be the cause of lymphopenia and overall immunosuppression, manifesting in this study as a profound insensitivity to Con A-induced liver inflammation. To gain further insight into the origin of the imbalance in T_eff_:T_reg_ ratio, we carried out a survey of T_reg_ frequency in the thymus and spleen.

**Figure 3.**
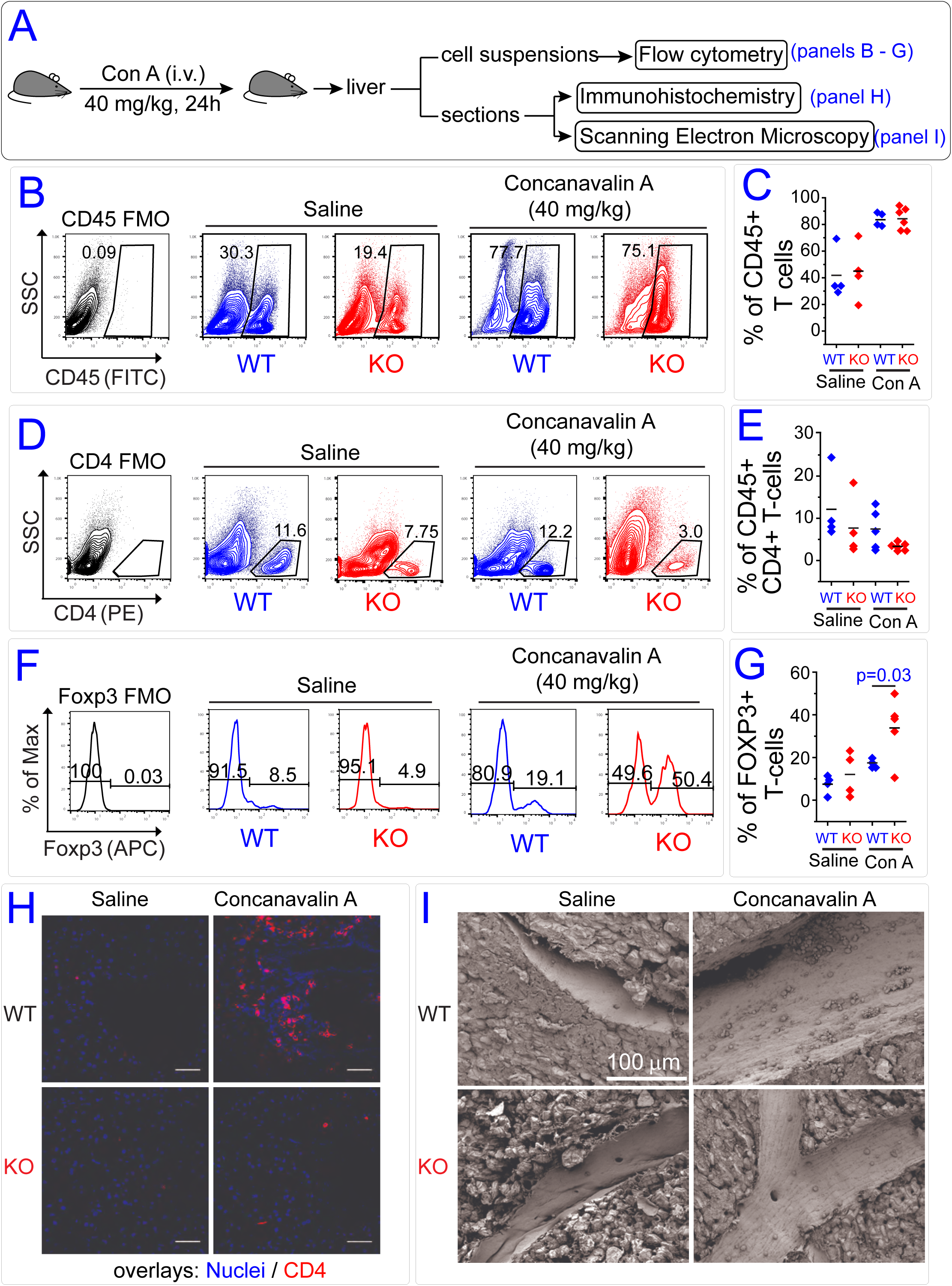
Increased infiltration of regulatory T cells in the livers of Con A-treated *Trpm7*^*fl*^ (*Lck Cre*) mice. **(A)** Experimental scheme. **(B)** Bivarant cytographs show infiltration of CD45+ hematopoietic cells in livers of *Trpm7*^*fl/fl*^ (WT) and *Trpm7*^*fl/fl*^*(LckCre)* (KO) mice, 24h after the administration of saline or Concanavalin A (40 mg/kg, iv). CD45 FMO refers to “<u>f</u>luorescence <u>m</u>inus <u>o</u>ne” control wherein cells were analyzed without staining with anti-CD45 antibody enabling accurate gating. **(C)** Scatter plot shows quantification of CD45+ hematopoietic cells based on flow cytometry analysis shown in *panel B*. Mean percentage of CD45+ cell in liver infiltrates are indicated by a dash. The data were collected from 4 independent experiments. **(D)** Bivarant cytographs showing infiltration of CD4+ T cells in livers of WT and KO mice, 24h after administration of saline or Con A (40 mg/kg, iv). **(E)** Scatter plot shows quantification of CD4+ T cells based on flow cytometry analysis shown in *panel D*. Mean percentages of CD45+ cells that are CD4+ are indicated by a dash. The data were collected from 4 independent experiments. **(F)** Flow cytomtery histograms of CD4+ cells expressing Foxp3 in the livers of WT and KO cells, 24h after administration of saline or Con A. The Foxp3 histograms were obtained from a cell population determined by sequential gating for a FSC/SSC profile, CD45+ and CD4+ (gating scheme shown in figure S3). **(G)** Scatter plots based on flow cytometry analysis (*panel F*) show quantification of Foxp3+ regulatory T cells in the livers of WT and KO mice. The p-value was calculated using t-test. The data were collected from 4 independent experiments. **(H)** Representative confocal microscopy images of indicated liver sections stained with anti-CD4 (red) and nuclear stain (blue). Scale bar represents 50 μm. **(I)** Representative Scanning Electron Microscope (SEM) images of indicated liver sections at 850X magnification. Note the reduced adhesion of immune cells in the endothelium of KO mice, relative to WT mice. Scale bar represents 100 μm.

### *Trpm7*^*fl*^*(Lck Cre)* mice have a higher frequency of CD4+, Foxp3+ regulatory T cells in thymus and spleen

We showed previously that the deletion of *Trpm7* in the T-cell lineage results in impaired T-cell development and a partial block in the transition from the double negative (CD4-CD8-, DN) to double positive (CD4+CD8+, DP) stage of T-cell development (Jin et al., 2008). This results in an accumulation of DN thymocytes and reduces the development of single positive (SP) CD4+ and CD8+ mature T cells (Jin et al., 2008). The SP cells egress to the periphery and populate all lymphoid organs but exhibit T-cell lymphopenia despite being resistant to apoptosis (Desai et al., 2012). As schematized **(Fig. 4A)**, we measured the frequency of T_reg_ cells in the thymus and spleen using flow cytometry. To assess the thymic development of FOXP3+ T_reg_ cells, we first gated on viable Thy1.^2+^ cells, selected CD8-CD4+ single positive mature T cells and then quantified the percentage of FOXP3+ T_reg_ cells in that population **(Fig. 4B)**. Consistently, we found that the thymi from the KO mice contained a higher frequency of FOXP3+ T_reg_ cells. As shown in the statistical box charts **(Fig. 4C)**, the thymi of the KO mice show a striking ∼3-fold increase in the frequency of FOXP3+ T_reg_ cells, when compared to WT thymi (WT 3.3±0.68%, n=8; KO 12.5±2.7%, n=7; p=0.004). Since the overall cellularity of thymi is significantly lower in the KO mice (Jin et al., 2008), the absolute numbers of T_reg_ cells in the KO thymi is comparable to WT thymi **(Fig. 4D)**. Importantly, this means that the KO thymi are exporting a T-cell composition that drastically overrepresents the T_reg_ population. The CD4+ thymocytes from KO mice also showed increased <u>M</u>edian <u>F</u>luorescence <u>I</u>ntensities (MFI) for FOXP3 staining **(Fig. 4E)**, indicating increased expression of the FOXP3 protein in cells. To check whether such a high frequency of T_reg_ cells is maintained in the periphery, we evaluated the composition of T-cell populations in the spleens. Consistently, we found that the CD4+ T cells in the KO spleens are composed of substantially higher percentage of FOXP3+ cells, when compared to WT spleens **(Fig. 4F)**. As shown in the box charts **(Fig. 4G)**, the mean percentage of splenic T_reg_ cells was more than 2-fold higher in KO mice (WT 11±1.5%, n=8; KO 24.5±1.7%, n=7; p=0.00005). Since the spleens of the KO mice contain a lower number of T cells (Jin et al., 2008), the mean absolute number of splenic T_reg_ cells in the KO mice was nearly identical to that quantified in WT mice **(Fig. 4H)**. Consistent with the CD4+ thymocytes, the CD4+ splenic T cells from the KO mice showed increased MFI for FOXP3 staining **(Fig. 4I)**, indicating increased cellular levels of the FOXP3 protein. We also analyzed the data by stipulating that T_reg_ cells be identified as Thy1.^2+^CD4+CD25+Foxp3+ cells and T_eff_ cells as Thy1.^2+^CD4+ cells. The representative contour plots and box charts show that the percentage of T_reg_ cells phenotyped in this manner are also significantly higher in thymi **(Fig. S4B)** and in spleens **(Fig. S4C)** of KO mice. With this analytical approach, the ratio of T_reg_:T_eff_ cells in the KO thymus trends higher than in WT mice – but is not significantly different. However, the Treg:Teff ratio is still significantly higher in the spleens of the KO mice, when compared to WT spleens **(Fig. S4D).** Thus, regardless of how T_reg_ cells are immunophenotyped, deletion of *Trpm7* increases the T_reg_:T_eff_ ratio significantly in the periphery. A very small percentage of CD8+ T cells is known to express FOXP3 (Churlaud et al., 2015) and we therefore checked if the KO mice also showed an abnormally high percentage of CD8+FOXP3+ T cells. As shown **(Fig. S4A)**, the percentage of FOXP3+CD8+ cells was very small and statistically identical in the WT and KO mice. To rule out a trivial explanation we checked whether the profound increase of T_reg_ frequency in KO mice was due to inefficient deletion of *Trpm7* in T_reg_ cells, giving them an undefined proliferative advantage over T_eff_ cells. To quantify the deletion of *Trpm7* in KO T_reg_ cells using freshly isolated CD4+CD25+ T_reg_ cells, we measured the relative mRNA levels of *Trpm7* that contain loxP-flanked *exon 17* and confirmed its efficient deletion in KO T_regs_ **(Fig. S4E)**. We also confirmed the loss of TRPM7 currents in KO Treg cells using patch clamp electrophysiology. To our knowledge, this is also the first patch-clamp recording of mouse T_reg_ cells and the recording conditions established to patch mouse T_reg_ cells are illustrated **(Fig. S4F)**. The WT CD4+CD25+ T_reg_ cells readily exhibited the characteristic outwardly rectifying ITRPM7 with a reversal potential at 0 mV (*left panel,* **Fig. 4J)**. This current was absent in KO T_regs_ and only a modest leak current was evident; this is also shown statistically in the box chart of current densities recorded at 100 mV, 5m after gaining electrical access to the cell (*right panel,* **Fig. 4J)**. These results clearly demonstrate that *Trpm7* is deleted efficiently in T_reg_ cells. Similar recordings also show that when compared to WT CD4+CD25-T_eff_ cells, the WT CD4+CD25+ T_reg_ cells ellicit significantly higher TRPM7 current densities **(Fig. 4K)**; the characteristic TRPM7 current-voltage relationship in these cells is shown **(Fig. S4G)**. The functional significance of increased TRPM7 currents in T_reg_ cells, in comparision to T_eff_ cells, is not yet clear. Next, we confirmed that the functional characteristics of *Trpm7-/-* T_reg_ cells are identical to that of WT T_reg_ cells.

**Figure 4.**
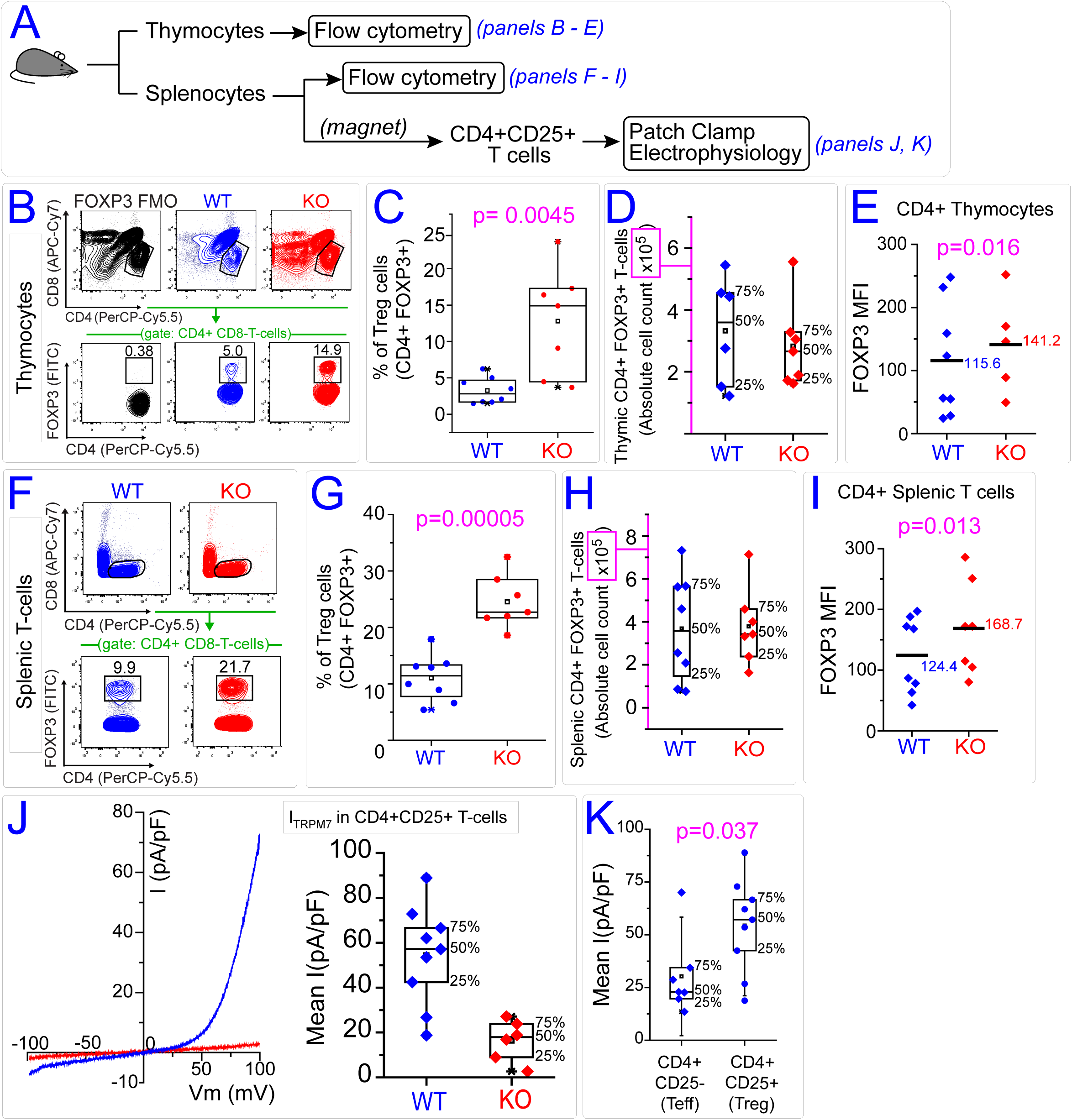
*Trpm7*^*fl*^ *(Lck Cre)* mice have a higher percentage of CD4+, Foxp3+ regulatory T cells in thymus and spleen. **(A)** Experimental scheme. **(B)** Flow cytometry contour plots sequentially gated on CD4+ and Foxp3+ regulatory T cells in *Trpm7*^*fl/fl*^(WT) and *Trpm7*^*fl/fl*^*(LckCre)* (KO) thymi. **(C)** Box charts show percentages of CD4+Foxp3+ T_reg_ cells in WT and KO thymi. The mean value is denoted by an empty square symbol and the median by a horizontal line. Other statistical features shown in the box charts are described in the methods. The p-value was calculated using two-tailed unpaired student t-test. The filled circles represent data obtained from individual mice. **(D)** Box charts show absolute numbers of CD4+Foxp3+ thymocytes in WT and KO thymi. **(E)** A scatter plot showing median fluorescence intensities (MFI) of FOXP3 on CD4+ thymocytes. **(F)** Flow cytometry contour plots sequentially gated on CD4+ and Foxp3+ regulatory T cells in WT and KO spleen. **(G)** Box charts show percentages of CD4+ Foxp3+ regulatory T cells in WT and KO spleen. **(H)** Box charts show absolute numbers of CD4+ Foxp3+ regulatory T cells in WT and KO spleen. **(I)** A scatter plot showing median fluorescence intensities (MFI) of FOXP3 on CD4+ splenic T cells. **(J)** I-V relationship (blue trace) of I_TRPM7_ in WT CD4+CD25+ (T_reg_) cells obtained by whole cell patch clamp recordings. Red trace confirms the absence of ITRPM7 in T_reg_ cells isolated from *Trpm7*^*fl*^ *(Lck Cre)* mice (KO). The patch clamp configuration and the recording conditions are shown in *figure S4F*. The box chart shows the I_TRPM7_ current densities quantified 5m after *break-in* at 100 mV in WT and KO T_reg_ cells. See methods for statistical parameters shown in the box chart. **(K)** The box chart shows the I_TRPM7_ current densities quantified 5m after break-in at 100 mV in Teff (CD4+CD25-) and T_reg_ (CD4+CD25+) cells isolated from WT mice. The p value was calculated using two-tailed unpaired student t-test and each filled circle represents data obtained from an individual cell.

### *Trpm7-/-* T_reg_ cells show normal cell surface and functional characteristics

Immunosuppression by T_reg_ cells is mediated by multiple mechanisms including inhibitory cytokines (IL-10, TGFβ), cytolysis of T_eff_ cells (Granzyme B), disruption of purinergic signaling by ectonucleotidases (CD39 and CD73) and inhibition of dendritic cell maturation (CTLA4) (Vignali et al., 2008). First, we evaluated their functional competency in *ex vivo* suppression assays wherein the proliferation of WT CD4+CD25-T_eff_ cells is suppressed by the presence of either WT or KO CD4+CD25+ T_reg_ cells, cocultured at varying ratios; this experimental design is illustrated **(Fig. 5A)**. The number of CFSE-labeled WT T_eff_ cells was held constant and the indicated ratios were achieved by varying the number of T_reg_ cells. The cells were then activated by a mixture of anti-CD3 and anti-CD28 for 3d and the cell division of CFSE-labeled T_eff_ cells was measured using flow cytometry. The histograms were analyzed to quantify the percentage of T_eff_ cells that underwent cell division **(Fig. 5B)**. Both WT and KO T_reg_ cells suppressed the proliferation of WT T_eff_ cells comparably. Next, we compared the cell surface density and gene expression of the key functional proteins CD39, CD73 and CTLA4 to determine the immunophenotypic features of KO T_reg_ cells in comparison to WT T_reg_ cells. We found that CD39 is expressed at moderate levels in CD4+FOXP3+ thymocytes but its expression levels increase in peripheral splenic T_reg_ cells. In the KO mice, this trend was preserved but we observed a substantially higher percentage of CD39+ T_reg_ cells in the thymus and spleen in comparision to WT T_reg_ cells **(Fig. 5C)**. A similar result was seen in the case of CD73 and CTLA4. As shown in the bar graphs **(Fig. 5D)**, the mean percentage *(left)* and MFI *(right)* of CD39+, CD73 and CTLA4 in thymi and spleens are significantly higher in KO than in WT mice. Through qPCR analysis, we also show that the gene expression of Interleukin 10 (*Il10*) and Granzyme B (*Gzmb*) was significantly higher in KO T_reg_ cells **(Fig. S5)**. The mRNA levels of *Ctla4*, *Lag3*, *Nt5e* and *Entpd1* were not statistically different but *Tgfb1* was lower in KO T_reg_ cells. Based on these data, we conclude that the overall expression of genes in KO T_reg_ cells is comparable but not identical to WT T_reg_ cells. The stability of FOXP3 expression is an important characteristic of T_reg_ cells, and it is thought that the thymically derived T_reg_ cells, also referred to as natural T_reg_ cells, exhibit increased FOXP3 stability (Sakaguchi et al., 2013). The thymus-derived natural T_reg_ cells can be identified on the basis of cell surface expression of Neuropilin-1 (Nrp-1) since this marker is expressed only at very low levels in peripherally induced T_reg_ cells (Weiss et al., 2012; Yadav et al., 2012). We found a substantially higher percentage of NRP-1+ KO T_reg_ cells in the thymus and spleen **(Fig. 5E** and **5F)**. These results indicate that the KO T_reg_ cells found in the periphery are derived through thymic differentiation and stably express *Foxp3*. In conjunction with the analysis of T_reg_-specific cell surface markers, gene expression and functional competency, we conclude that the TRPM7-/-T_reg_ cells are functional and because of their increased frequency, contribute resistance to experimental hepatitis.

**Figure 5.**
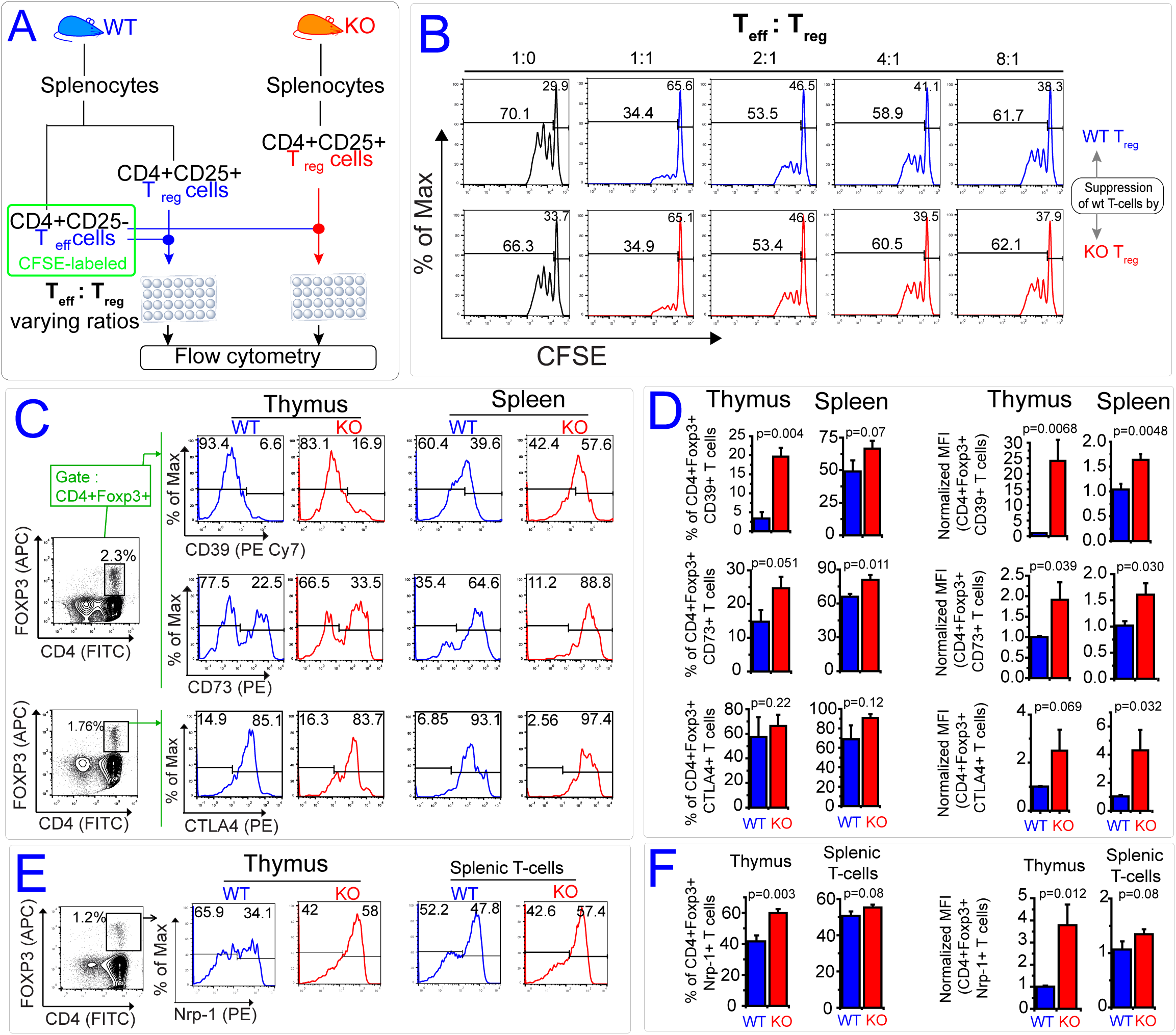
Characterization of *Trpm7-/-* T_reg_ cells isolated from the thymus and lymph nodes of *Trpm7*^*fl*^ *(Lck Cre)* mice. **(A)** The experimental scheme used for T_reg_ suppression assay. **(B)** Suppression of CFSE-labeled wt T-cell proliferation after 3 days of culture in the presence of T_reg_ cells isolated from WT and KO mice. The indicated ratios indicate the proportion of effector T cells (CD4+CD25-* T_eff_) relative to regulatory T cells (CD4+CD25+, T_reg_) in the cultures. Histograms show the dilution of CFSE in T_eff_ cells as measured by flow cytomtery. Percentage of T_eff_ cells showing dilution of CFSE dye reflects the percentage of T_eff_ cells that have undergone cell division. **(C)** T_reg_ cells from the thymi and spleens of *Trpm7*^*fl/fl*^ (WT) and *Trpm7*^*fl/fl*^*(LckCre)* (KO) mice were identified by flow cytometry (see bivariant gating of CD4 and Foxp3 shown on the left) and further analyzed for the expression of T_reg_ functional markers CD39, CD73 and CTLA4 (histograms). **(D)** The bar graphs shows the percentage (left) MFI (right) of CD39, CD73 and CTLA4 in thymus and spleen of WT and KO mice. The data were collected from 2 independent experiments with a total sample size of 4 mice. The p-value was calculated by using one–tailed unpaired student t-test. **(E)** Cell surface expression of neuropilin-1 (Nrp-1) on thymic and splenic T_reg_ cells (CD4+ Foxp3+) from WT and KO mice. **(F)** The bar graphs shows the percentage (left) MFI (right) of Nrp-1 in thymus and splenic T-cells of WT and KO mice. The data were collected from 2 independent experiments with a total sample size of 4 mice. The p-value was calculated by using one–tailed unpaired student t-test.

### Deletion of Trpm7 in thymocytes and T cells increases IL-2 sensitivity

IL-2 plays a critical role in regulating the development, maintenance and function of T_reg_ cells (Bayer et al., 2013; Fontenot et al., 2005; Malek, 2003) through the activity of STAT5, a key transcription factor responsible for *Foxp3* gene expression (Antov et al., 2003; Burchill et al., 2007). To assess the state of IL-2-signaling in the developing T_reg_ cells by flow cytometry, we first checked IL-2Rα 2CD252 mRNA and protein in the KO thymocytes. The IL-2Rα mRNA expression in thymocytes is five-fold higher in KO mice compared to WT **(Fig. 6A)** and accordingly protein expression is also increased in KO thymocytes **(Fig. 6B).** Interestingly, deletion of *Trpm7* also increases the output of IL-2 by the KO thymocytes. Analysis of intracellular IL-2 in Thy1.^2+^ thymocytes shows that KO thymocytes register a significant increase in the MFI of intracellular IL-2 staining – the histogram **(Fig. 6C)** and quantification **(Fig. 6D)** are shown. Similar differences in IL-2 production are observed when the analysis is restricted to Thy1.^2+^CD4+ thymocytes **(Fig. 6E** and **6F)**. Although the *Trpm7*-/-thymocytes produce increased IL-2 in the thymus, the *Trpm7*-/-T-cells in the periphery do not produce significantly more IL-2 than the WT T-cells. The measurements were carried out in the Thy1.^2+^ CD4+ T-cells obtained from the spleen and the mesenteric lymph node and are shown **(Fig. S6A)**. The concentration of IL-2 in the serum of WT and KO mice is also identical **(Fig. S6B)**. These results show that the deletion of *Trpm7* in the T-cell lineage increases the expression of IL-2 and IL-2Rα in the developing thymocytes and suggest that this substantial increase in IL-2 promotes the development of FOXP3+ T_reg_ cells. To confirm that the IL-2 signaling is indeed potentiated, we evaluated the phosphorylation of STAT5, a transcription factor activated by IL-2Rα to induce FOXP3 expression in developing T_reg_ cells. The overlaid histograms for intracellular phospho-STAT5 (or pSTAT5) staining in thymocytes obtained from WT and KO thymi are shown **(Fig. 6G)**. The frequency of pSTAT5+ cells as well as the MFI of pSTAT5, a measure of STAT5 activity per cell, are significantly higher in KO thymocytes **(Fig. 6H)**. The increased pSTAT5 was confined to thymic T_reg_ cells; freshly isolated splenic T cells show no difference in pSTAT5 staining **(Fig. S6F)**. Collectively, these data indicate that the thymocytes in KO mice show increased IL-2-signaling, the major cytokine driving STAT5 activation. Since IL-2Rα is also a transcriptional target of STAT5, we reasoned that *Trpm7*-/-thymocytes maintain high IL-2Rα levels on the surface and increased STAT5 activity through a feed-forward positive feedback loop. This idea is supported by the measurement of the occupancy of STAT5 on the IL-2Rα promoter using chromatin immunoprecipitation coupled to qPCR (ChIP-qPCR). Immunoprecipitation of the cross-linked chromatic was carried out using a ChIP-grade anti-pSTAT5 or control IgGα antibody and the bound genomic DNA was probed by qPCR using primers designed to amplify the IL-2Rα promoter sequences. We found a 2-fold enrichment of IL-2Rα promoter sequence in anti-pSTAT5 chromatin IPs from KO thymocytes, when compared to WT thymocytes **(Fig. S6G)**. To further substantiate this model, we hypothesized that TRPM7 restrains the JAK-STAT pathway downstream of IL-2 receptor and tested the prediction that *Trpm7-/-* thymocytes treated with IL-2 *ex vivo* would also exhibit higher activation of STAT5. As shown by the bivariant plots for pSTAT5 and FOXP3 **(Fig. 6I;** gating strategy in **Figure S6C)**, when WT thymocytes are treated with IL-2, a very small percentage of cells exhibit phosphorylation of STAT5 at 5m or at 30m. In contrast, when KO thymocytes are treated with IL-2, the percentage of pSTAT5+FOXP3+ cells is substantially higher (upper right quadrant). The bar graph shown in **Figure 6J**, clearly shows a nearly 5-fold increase in the percentage of pSTAT5+FOXP3+ cells 30m after treatment with 100 ng/ml IL-2 (n=4; WT 0.58±0.15%; KO 2.61±1.4%; p=0.014). We also quantified the percentage of STAT5+ thymocytes and found a similar increase in KO thymocytes **(Fig. S6D)**. These results argue that the increased IL-2 sensitivity of KO thymocytes is an intrinsic property of *Trpm7*-/-thymocytes and not limited to differentiated T_reg_ cells alone. Although we did not see a difference in basal activation of STAT5 in freshly isolated splenic T cells, we probed the sensitivity to *ex vivo* IL-2 stimulation. The splenic T cells from KO mice respond rapidly to IL-2 stimulation by phosphorylating STAT5 and show a nearly 3-fold increase in the percentage of pSTAT5+FOXP3+ cells when compared to WT mice **(Figures 6K** and **6L)**. As shown **(Fig. S6E)**, the increased IL-2-sensitivity seen in KO T cells is not limited to FOXP3+ cells. The MFI of phospho-STAT5 staining was significantly higher in IL-2 treated CD4+FOXP3+ T cells from the KO mice **(Fig. S6H)**, supporting the conclusion that the absence of TRPM7 increases IL-2 sensitivity of T-cells. Overall, these findings clearly demonstrate that the deletion of *Trpm7* potentiates the development of T_reg_ cells by sensitizing the thymocytes to IL-2 stimulation and promoting the activation of STAT5, a key transcription factor for *Foxp3* transcriptional regulation. Based on this model we evaluated whether the deletion of *Trpm7* or pharmacological inhibition of TRPM7 promotes the transcription of *Foxp3* in IL-2 activated thymocytes *ex vivo*.

**Figure 6.**
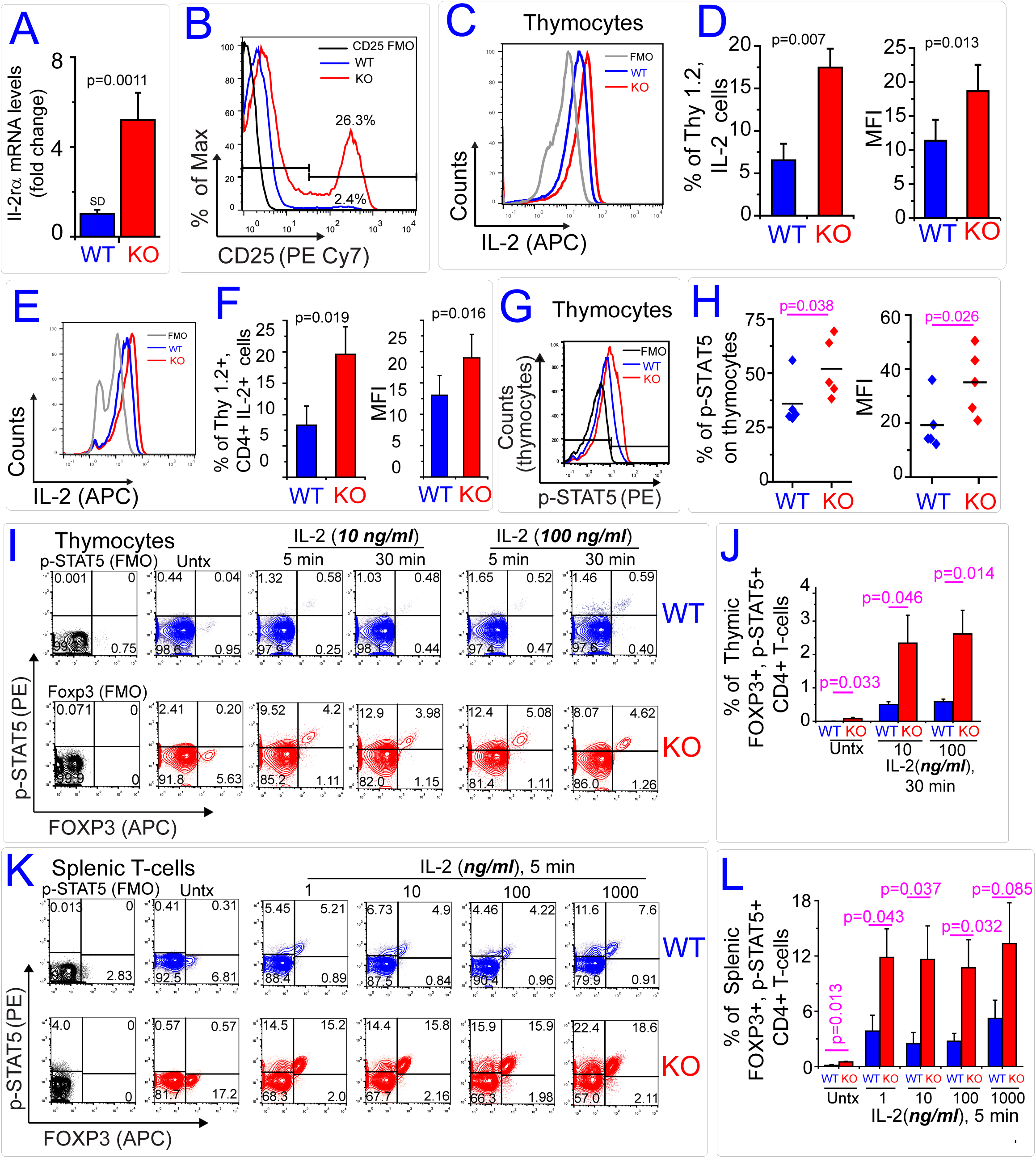
*Trpm7-/-* thymocytes and T cells show increased phosphorylation of STAT5 in response to IL-2 (A) RT-qPCR analyses of Il-2Rα mRNA expression in WT and KO thymocytes. The relative fold change was calculated with internal control (β2m). The p value was calculated using two–tailed unpaired student t-test (n=3). **(B)** Flow cytometry histograms show the cell surface expression of Il-2Rα 2CD25) in WT and KO thymocytes. The data are representative of 7 independent experiments. **(C)** Overlaid flow cytometry histograms showing the intracellular IL-2 cytokine in thymocytes gated on Thy 1.2+ cells. The cells were analyzed after isolation from WT (blue) and KO (red). The gray line is an FMO control. **(D)** The bar graph on the left denotes the percentage of IL-1.2+Thy1. 1.2+ thymocytes obtained from WT (blue) and KO (red) mice. The bar graph on the right shows MFI of IL-2 staining in Thy1. 1.2+ thymocytes. **(E)** Overlaid flow cytometry histograms show the staining of intracellular IL-2 in Thy1. 1.2+ CD4+ thymocytes obtained from WT and KO mice. **(F)** The bar graph denotes the average percentage of IL-1.2+ CD4+Thy1.2+ thymocytes in WT and KO mice and the error bars represent SEM (left). The bar graph on the right shows the MFI of IL-2 staining in CD4+Th1. 1.2+ thymocytes and the error bars represent SEM. The data were collected from 3 independent experiments (n= 5 mice) and the p values were calculated using unpaired student t-test. **(G)** Flow cytometry histogram shows intracellular staining of phospho-STAT5 (pSTAT5) in thymocytes freshly isolated from WT and KO mice. *<u>F</u>luorescence <u>m</u>inus <u>o</u>ne* (FMO) control reflects background staining in WT thymocytes without using anti-pSTAT5 antibody. **(H)** Scatter chart on the left shows percentage of pSTAT5+ thymocytes, freshly isolated from WT and KO mice using the flow cytometry analysis shown in *panel G*. Scatter chart on the right shows the <u>m</u>edian <u>f</u>luorescent <u>i</u>ntensities (MFI) of pSTAT5 staining, a measurement of pSTAT5 protein levels, in WT and KO thymocytes. The p values were calculated using two–tailed unpaired student t-test. **(I)** Bivariant contour plots show Foxp3 and pSTAT5 staining of CD4+ thymocytes from WT and KO mice, before and after IL-2 stimulation. The IL2 concentrations and time of IL-2 treatment are indicated. Analysis is confined to previously gated CD4+ thymocytes; gating strategy is shown in *figure S6C*. **(J)** Bar graph shows the percentages of pSTAT5+ T_reg_ cells as determined by upper right quadrant gate from flow cytomtery analysis shown in *panel I*. The error bars reflect SEM (n=4); p-values calculated using t-test **(K)** Bivariant contour plots show Foxp3 and pSTAT5 staining of CD4+ splenic T cells obtained from WT and KO mice, before and after 5 min of IL-2 stimulation (with indicated concentrations). Analysis is confined to previously gated CD4+ cells. See *figure S6C* for gating. **(L)** Bar graph shows the percentages of pSTAT5+ splenic T_reg_ cells as determined by upper right quadrant gate from flow cytomtery analysis shown in *panel K*. Error bars reflect SEM (n=4); p-values calculated using t-test.

### Deletion or pharmacological inhibition of TRPM7 enhances the induction of *Foxp3* during *ex vivo* activation of thymocytes

When WT thymocytes are activated *ex vivo* by anti-CD3 antibodies in the presence of IL-2, a modest upregulation of *Foxp3* transcripts is evident by qRT-PCR. The deletion of *Trpm7* increases *Foxp3* transcription (*left panel***, Fig. 7A)** but this increased *Foxp3* transcription in KO thymocytes is abrogated in the presence of a highly selective JAK3 inhibitor (CP690550, 2 nM) (*right panel***, Fig. 7A)**. We activated the thymocytes *ex vivo* and evaluated the differentiation of T_reg_ cells (CD4+CD25+FOXP3+) after 48h. Activation by anti-CD3 antibodies and IL-2 resulted in a modest increase in the percentage of WT CD4+FOXP3+ thymocytes but the percentage of KO CD4+FOXP3+ thymocytes was significantly higher **(Fig. 7B)**. The bar charts in **Figure 7C** show that when activated in presence of IL-2, the KO thymocytes readily differentiate toward the T_reg_ lineage. The increased differentiation of CD4+FOXP3+ in presence of anti-CD3 alone is likely due to autocrine production of IL-2 that activates JAK3 and STAT5 through the common γ chain. Since TRPM7 is an ion channel and a serine-threonine kinase domain, we investigated whether the pharmacological inhibition of TRPM7 channel is sufficient to recapitulate the effects of *Trpm7* deletion. To evaluate the role of the TRPM7 channel specifically, we used FTY720, an FDA-approved Sphingosine-1-Phosphate Receptor (S1PR)-targeting prodrug that is phosphorylated *in vivo* to FTY720-P, an analog of S1P (Brinkmann et al., 2010). FTY720 blocks TRPM7 channel without further modification to its phosphorylated form (Qin et al., 2013). Since the previous report identified FTY720 as a channel blocker of TRPM7 using ectopic expression of TRPM7 in cell lines (Qin et al., 2013), we first confirmed that FTY720 potently blocks native I_TRPM7_ in mouse thymocytes. The inhibition of the characteristic outwardly rectifying TRPM7 current in thymocytes by 2 μM FTY720 is shown **(Fig. 7D)**. The residual current seen in the presence of FTY720 includes leak currents indicating that the inhibition shown in the inset bar graph is an underestimate of TRPM7 inhibition. The phosphorylated derivative FTY720-P binds to S1PR, but does not block TRPM7 - this is illustrated **(Fig. 7E)**. We activated WT thymocytes in the presence of FTY720 and FTY720-P to assess the effect on *Foxp3* gene expression. In the presence of FTY720, but not in the presence of FTY720-P, the thymocytes showed a potent upregulation of Foxp3 mRNA **(Fig. 7F)**. Since FTY720-P, but not FTY720, is the agonist of S1PR, these results clearly indicate that the increased transcriptional _reg_ulation of *Foxp3* is not mediated by S1PR. Using an identical experimental setup, we then measured the protein expression of FOXP3 in the presence of FTY720. Inhibition of TRPM7 by FTY720 greatly increased the expression of FOXP3 in thymocytes that are activated *ex vivo* **(Fig. 7G)**, with a concomitant increase in STAT5 phosphorylation **(Figures 7H** and **7I)**. We also investigated whether administration of FTY720 (i.p.) can increase T_reg_ cells in WT mice. We administered FTY720 into WT and KO mice (3 doses for 6 days at 0.1μg/kg) and measured CD4+FOXP3+ T cells **(Fig. S7C** and **S7D)**. Wild type mice displayed a ∼50% increase of percentage of T_reg_ in the thymus, compared to saline treated mice. However, FTY720 had no significant increase in the percentage of T_regs_ in KO mice, compared to saline treated KO mice **(Fig. S7D)**. Overall, these data bolster the framework that TRPM7 channel is a negative regulator of JAK-STAT signaling downstream of IL-2 receptor. In *Trpm7-/-* thymocytes or when TRPM7 channel is blocked in WT thymocytes, the activation of STAT5 is potentiated, resulting in increased transcription of *Foxp3* and increased thymic differentiation of CD4+FOXP3+ Treg cells. Since IL-2-dependent STAT5 activation also plays a role in the induction of peripheral T_reg_ cells, we evaluated the *ex vivo* induction of splenic CD4+CD25-cells in the presence of TGF-β1 and IL-2. We found that *Trpm7-/-* T cells show modestly increased propensity to differentiate as CD4+CD25+FOXP3+ T_reg_ cells **(Figures S7A** and **S7B)**. These results are in harmony with our proposed model that TRPM7 restrains IL-2 mediated STAT5 activation and subsequent *Foxp3* gene expression. A graphical summary of our findings in the context of the proposed model is shown in **Figure S7E**.

**Figure 7.**
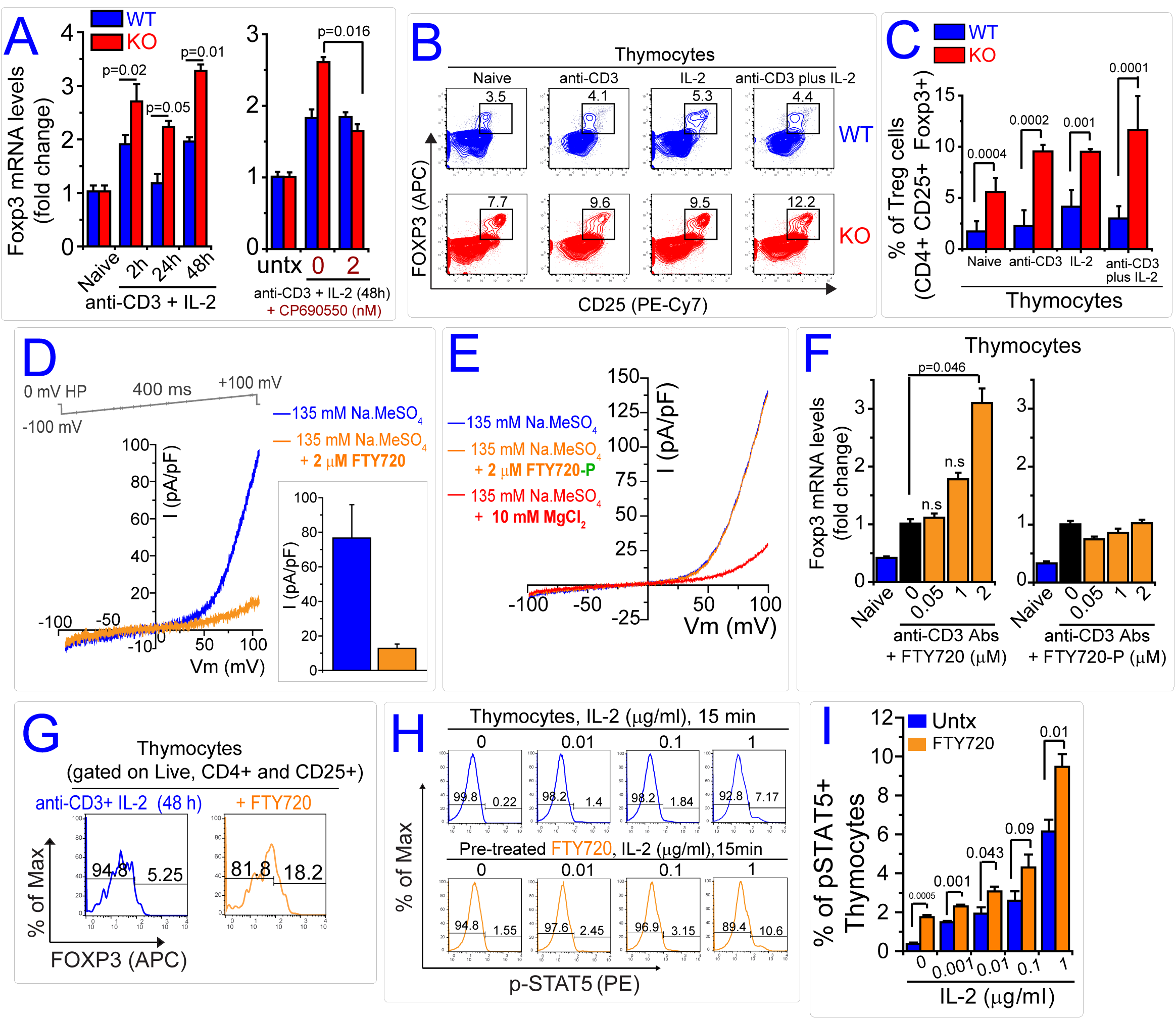
*Foxp3* gene induction is enhanced by deletion or inhibition of TRPM7. **(A)** qRT-PCR showing the upregulation of *Foxp3* expression in WT and KO thymocytes activated with anti-CD3 + IL-2 at different time points prior to analysis. The right panel shows the suppression of Foxp3 induction in KO thymocytes in presence of 2 nM JAK inhibitor CP690550. Both bar graphs denote mean values and the error bars reflect SEM (n=3). The p value was calculated using student’s t-test. **(B)** Bivariant contour plots showing the analysis of Foxp3 and CD25 expression on previously gated CD4+ thymocytes after activation in indicated conditions. **(C)** Bar graph shows quantification of the percentages of CD4+CD25+Foxp3+ T_reg_ cells using the analysis shown in *panel B*. Error bars reflect SEM (wt, n= 3 and ko, n=4). Each experiment used pooled thymocytes from at least 2 mice. The p value was calculated using two-tailed unpaired student’s t-test. **(D)** I-V relationship (blue trace) of I_TRPM7_ in WT thymocytes obtained by whole cell patch clamp recordings. Orange trace shows the inhibition of I_TRPM7_ after bath perfusion of 2 μM FTY720. Inset bar graph shows I_TRPM7_ current densities in thymocytes obtained 5m after break-in before and after perfusion with 2 μMFTY720 at 100 mV (n=3). **(E)** I-V relationship of I_TRPM7_ in response to bath perfusion of S1PR agonist FTY720-phosphate (2 μM) shows that FTY720-P does not block TRPM7. The orange trace (FTY720-P) overlaps with the blue trace (without FTY720-P), masking it. Inhibition of I_TRPM7_ by MgCl_2_ is shown (red trace). **(F)** qRT-PCR based quantification of Foxp3 levels in WT thymcytes activated with anti-CD3 ab in presence of FTY720 (left panel) or FTY720P (right panel) at indicated concentrations (48 hours). Bar graphs reflect SEM (n=3). The p value, calculated using unpaired student t-test is denoted on the bar graph. **(G)** Flow cytometry histogram shows intracellular staining of Foxp3 on live CD4+CD25+ thymocytes isolated from WT mice. Before analysis, WT thymocytes were activated for 2 days with anti-CD3 and IL-2, in absence (blue) or in presence of FTY720 (orange). **(H)** Flow cytometry histogram of intracellular phospho-STAT5 staining. Thymocytes were pretreated with FTY720 (orange) or left untreated (blue) and then stimulated with IL-2 at indicated concentrations for 15 min. **(I)** Bar graph quantifies the percentages of pSTAT5+ thymocytes from flow cytometry analysis shown in *panel H*. The data are representative of 3 independent experiments where the cells were cultured in triplicates. The denoted p value was calculated using student’s t-test.

## DISCUSSION

In the clinic, adoptive transfer of T_reg_ cells has been shown to alleviate a wide variety of autoimmune diseases (Trzonkowski et al., 2015). However, these efforts face major technical hurdles due to insufficient knowledge of how T_reg_ cells develop, survive and function (Trzonkowski et al., 2015). The manipulation of T_reg_ numbers through pharmacological approaches may be a far more attractive immunomodulatory therapy for autoimmune diseases, cancer and organ transplantation. Despite the potential clinical impact of such T_reg_-targeted pharmacology, there is a major dearth of “druggable” molecular targets capable of manipulating T_reg_ development and frequency. Our study addresses this salient gap by identifying TRPM7 as a key novel controller of T_reg_ development. We demonstrate that the genetic deletion of *Trpm7* or pharmacological inhibition of TRPM7 channel promotes the development of functional T_reg_ cells. Thus, TRPM7 is the first ion channel target shown to increase T_reg_ numbers when blocked, and these findings reveal a novel drug development path toward the therapeutic manipulation of T_reg_ numbers.

Our study shows that TRPM7 channel activity restrains IL-2 dependent activation of STAT5. IL-2 plays a critical role in immune tolerance by regulating the development, maintenance and function of T_reg_ cells (Bayer et al., 2013; Fontenot et al., 2005; Malek, 2003). Although it acts as a growth factor for activated T cells, deletion of IL-2 and IL-2 receptor components causes a lymphoproliferative disorder accompanied by autoimmune disease (Sadlack et al., 1993; Suzuki et al., 1995; Willerford et al., 1995). Remarkably, adoptive transfer of T_reg_ cells alone prevents the lethal autoimmunity in IL-2Rβ-deficient mice demonstrating that the key function of IL-2 signaling is intricately tied to the normal development and maintenance of functional T_reg_ cells (Malek et al., 2002). When IL-2 signaling is impaired, FOXP3+ T_reg_ cells develop in the thymus but they express low levels of FOXP3 and show functional deficiencies due to reduced expression of CD39, CD73 and CTLA4 (Bayer et al., 2007; Cheng et al., 2013; D’Cruz and Klein, 2005; Fontenot et al., 2005). In the absence of IL-2 signaling, other common γ-chain cytokines appear to play a compensatory role to induce *Foxp3* expression but in the absence of γ-chain, the thymic development of FOXP3+ T_reg_ cells is completely abolished (Fontenot et al., 2005). Since the key transcription factor downstream of γ-chain-induced JAK-STAT signaling in thymocytes, is STAT5 (Antov et al., 2003; Burchill et al., 2007), enhanced IL-2 sensitivity underlies the mechanistic basis for increased T_reg_ development in the KO mice. Consistently, the increased cell surface levels of CD25, a STAT5 target gene (Kanai et al., 2014), likely contributes in a feed forward loop to increase IL-2 sensitivity in *Trpm7-/-* T cells. In the lymph nodes, IL-2 signaling and STAT5 mediate a crucial function in enabling the T_reg_ cells to constrain the local expansion of conventional T cells into damage-causing T_eff_ cells (Liu et al., 2015). Future studies will evaluate whether *Trpm7-/-* T_reg_ cells are hyper-vigilant in this aspect by virtue of their increased sensitivity to IL-2.

The mechanism through which TRPM7 channel activity restrains the JAK-STAT pathway remains unclear at this point and will be a major focus of our future studies. TRPM7 channel activity is polymodally regulated by changes in internal Mg^2+^(Kozak and Cahalan, 2003), phospholipids (Runnels et al., 2002), pH (Jiang et al., 2005) and C-terminal proteolysis (Desai et al., 2012; Krapivinsky et al., 2014). In physiological conditions, the influx of Ca^2+^, Na^+^, Zn^2+^and Mg^2+^through TRPM7 would result in membrane depolarization, and trigger signaling events sensitive to Ca^2+^and Zn^2+^. However, JAK-STAT pathway has not been previously reported to be regulated by such electrical signaling and at this point, no particular mode of ionic regulation can be ruled out. In the case of mice where store-operated Ca^2+^entry through Orai channels is abolished selectively in the T-cell lineage, the mice fail to develop an adequate number of T_reg_ cells (Oh-Hora et al., 2008). From the standpoint of the two-step model (Lio and Hsieh, 2008) of T_reg_ development, Ca^2+^entry through the Orai channels is likely essential for the strong TCR-dependent signals that drive the upregulation of IL-2 receptor and other components of IL-2 signaling pathway in the developing thymocytes. In contrast, the function of TRPM7 is to restrain the IL-2 mediated signals that finalizes the STAT5-dependent expression of *Foxp3.* In essence, the functions of Orai and TRPM7 channels are designed to be in opposition during the process of T_reg_ development. Hence the general design principles underlying the checks and balance in immune regulation appear to be preserved even at the scale of rapid electrical signaling (Germain, 2012).

In the context of S1PR pharmacology, FTY720 functions as a prodrug, which is phosphorylated *in vivo* to FTY720-P, the actual ligand for S1PR. Binding of S1PR by FTY720-P, but not by FTY720, promotes the sequestration of lymphocytes in lymph nodes, resulting in immunosuppression. Through patch clamp recordings of thymocytes, we confirmed that TRPM7 channel is blocked by FTY720, but not by FTY720-P; this enables the inhibition of TRPM7 without modulating S1PR. In presence of FTY720, but not FTY720-P, IL-2-stimulation of activated thymocytes increases the transcription of *Foxp3* and promotes the ex vivo differentiation of T_reg_ cells. This result shows that blocking TRPM7 channel activity promotes the differentiation of T_reg_ cells. Importantly, treating human patients with FTY720 has been reported to increase the frequency of T_reg_ cells (Muls et al., 2014) supporting the conclusion that at least some component of FTY720-mediated immunosuppression is mediated by increasing the frequency and function of T_reg_ cells through TRPM7 channel inhibition. Although the involvement of TRPM7 channel activity is clearly indicated by experiments involving FTY720, the role of the TRPM7 kinase domain cannot be ruled out. The cleavage of TRPM7 at D1510 increases the ion channel activity and liberates a functional kinase domain (M7CK-S) (Desai et al., 2012) while cleavage sites N-terminal of D1510 inactivate the channel and liberate a longer kinase domain (M7CK-L) that translocates to the nucleus and modifies the chromatin by phosphorylating the histones^23^. It remains possible that during T cell development, the kinase domain plays an important role in reprogramming the chromatin landscape.

The novel and significant insight derived from this study is the surprising function of the ion channel TRPM7 in controlling T_reg_ development and frequency. Since TRPM7 is an attractive drug target, and the lead compound FTY720 is already an FDA-approved drug, the findings have major implications for immediate translational research as well as medicinal chemistry approaches for more selective blockers of TRPM7. The mechanistic insights open the door toward a deeper investigation of TRPM7 and other ion channels in the development, function and stability of T_reg_ cells. In future, it may be possible to increase the T_reg_ numbers *in vivo* by TRPM7-targeting drugs and thereby induce tolerance in patients suffering from autoimmunity.

## EXPERIMENTAL PROCEDURES

*All procedures are described in further detail in Supplemental Experimental Procedures (SEP).*

### Mice

Mice were housed and bred in accordance with policies and protocols approved by the Institutional Animal Care and Use Committee (IACUC) of University of Virginia. Male and female mice aged between 4-12 weeks were used for the experiments.

### Reagents

Antibodies, purified cytokines, chemicals and commercial kits used in this study are described in SEP.

### Concanavalin A induced hepatitis model

Age and sex-matched WT and KO mice were weighed 2 hours prior to injection (i.v) of Concanavalin A (40 mg/kg) or saline (PBS) to determine appropriate dosage. Immediately after euthanasia (at indicated time points), blood and liver were collected for analysis in a blinded manner.

### Isolation of cells and flow cytometry

PBS-perfused livers were excised, washed in Isakov’s Modified Dulbecco’s Medium (with 10% FBS) and digested in 0.05% collagenase (37°C, 25 min). Mononuclear cells (MNCs) containing the immune cells were purified using a 40% Histodenz (Sigma-Aldrich) gradient after centrifugation (1500g, 20 min, 4°C). Spleens, thymi and lymph nodes were physically disrupted in cold, sterile RPMI-1640. Cell suspensions were filtered through 70 μm nylon mesh and contaminating erythrocytes were lysed using ACK buffer. The cell suspensions were then stained for flow cytometry or subjected to further magnetic separation to isolate different subsets using protocols provided by the manufacturer. For intracellular staining of Foxp3, the cells were fixed in 1% paraformaldehyde and permeabilized with 0.1% saponin. Intracellular staining of phospho-STAT5 (pSTAT5) was done after concomitant fixation and permeabilization with 100% methanol.

### T-cell activation, CFSE proliferation assay and transwell-migration assay

T cells were cultured with a cell density of 0.5 ×; 10^6^/ml in a culture volume of 200 μL/well in 96 well round bottom plates. The cells were activated with Concanavalin A (5 μg/ml, 48h) or as indicated. Alternatively, cells were activated using a mixture of anti-CD3ε (2.5 μg/ml) and anti-CD28 (1 μg/ml) for 48h. After activation, cells or the supernatants were harvested for analysis. For CFSE proliferation assays, the T-cell subsets were cultured at a reduced density (50000/well) and stimulated (for proliferation) using a mixture of anti-CD32_ε_ (0.25 μg/ml) and anti-CD28 (0.125 μg/ml). For transwell-migration assay, the migration of T cells from the upper chamber to the lower chamber, through the membrane (5.0 μm pore size membrane), in response to conditioned medium from LPS-stimulated BMDMs was analyzed. T cells that migrated to the bottom chamber were manually counted using trypan blue and the data are represented as a percentage of T cells that migrated across the membrane.

### Electrophysiology

TRPM7 currents (I_TRMP7_) were measured in the whole cell configuration as illustrated **(Fig S4F)**. T cells were activated with a mixture of anti-CD3ε and anti-CD28 antibody (or Concanavalin A) and cultured in 96-well plate (37°C, 5% CO_2_). FTY720 (2 μM) or MgCl_2_ (10 mM) were used with external solution to inhibit TRPM7 currents. All currents were recorded using an Axopatch 200B amplifier (Molecular devices, Sunnyvale, CA). The recording protocol consisted of 400 ms ramps from −100 mV to +100 mV and holding potential (HP) of 0 mV. The signals were low-pass filtered at 5 kHz and sampled at 10 kHz. All electrophysiology experiments were done at RT (∼22°C). The average current densities were plotted with the relevant statistical information as a box charts (see *Statistics* below).

### Suppression assay

Splenic cell suspensions were isolated from WT and KO mice. Effector T cells (T_eff_) or regulatory T cells (T_reg_) were then isolated through negative and positive magnetic selection using Dynabeads Flow Comp Mouse CD4+ CD25+ T_reg_ Cells kit (Lifetech). T_eff_ cells were labeled with 5μM CellTrace CFSE, washed and cultured in 96-well round bottom plate and stimulated with a mixture of anti-CD3ε and anti-CD28 antibodies (5% CO_2_, 37°C, 3 days). The number of T_eff_ cells was kept constant at 5 10^4^ cell per well, whereas the number of co-cultured T_reg_ cells varied in accordance with indicated ratios. For flow cytometry analysis, the cells were collected and stained with 7-AAD (to assess viability), anti-CD4 and anti-CD25 and the data are represented as the percentage of T_eff_ cells showing proliferation-induced dilution of CellTrace CFSE dye. This parameter reflects the percentage of cells undergoing cell division.

### *Ex vivo* generation of regulatory T cells from thymocytes

Thymocytes were isolated from WT and KO as single cell suspensions and cultured in X-vivo 15 growth medium (Sartorius Biotech, Cat #04-744Q) supplemented with 100 U/ml penicillin, and 100 μg/ml streptomycin. Thymocytes were activated with anti-CD3ε antibody in presence of recombinant hIL-2 (10 ng/ml) in 96-well round bottom plates (5% CO_2_, 37°C, 48 h). After 48h, thymocytes were collected and were either used to isolate mRNA for Foxp3 mRNA analysis by qPCR or analyzed by flow cytometry for immunophenotyping and intracellular Foxp3 expression.

### Statistics

Box charts were plotted using data analysis and graphing software Origin Pro 9.1.0 (Origin Lab). Statistical box charts in *figures 4C, 4G* and *S4B* are shown as Box (range of 25-75 percentile), whisker bars (1-99 percentile) and data overlap. Each data point including outliers is shown along with median (horizontal line) and arithmetic mean (empty square). Box charts in *figure 4D, 4H, 4K and 4J* are shown as box (range of percentile 25-75) and whisker bars (SD). All data points including outliers are shown with median (horizontal line) and mean (empty square) in the box. The p-values were calculated using two tailed unpaired student t-test and shown on each figure. Sample size equals the number of data points shown in the box chart.

## AUTHOR CONTRIBUTIONS

**Conceptualization:** EKM, SKM, BND; **Methodology:** SKM, MSS, EKM, TJB, BND; **Investigation:** SKM, MSS, BND, JKK, JSR, EJS, CAP; **Manuscript writing – Original Draft:** BND; **Manuscript writing – Review&Editing:** SKM, MSS, EKM, TJB, BND; **Supervision:** SKM, TJB, BND; **Funding Acquisition:** TJB, BND; **Project Administration:** BND

## ACKNOWLEDGEMENTS

We thank Marta Stremska, Phil Seegren and Kalina Szteyn for helpful discussions and suggestions. We also thank UVA flow cytometry core, UVA Advanced Microscopy Facility and UVA mouse care facilities for technical assistance. Lastly, we are very grateful for the following sources of funding to BND: NIH (GM108989, HL120840), American Cancer Society (UVA ACSIRG). Similarly, MSS is grateful for pre-doctoral training support from NIH (5T32GM007055-40, −41).

## SUPPLEMENTAL INFORMATION

## SUPPLEMENTAL EXPERIMENTAL PROCEDURES

### Animals

Male and female mice aged between 4-12 weeks were used for the experiments. *Trpm7*^*fl*^ mice wherein exon 17 of *Trpm7* is flanked by LoxP sites *and Trpm7*^*fl*^*(Lck Cre)* mice which deletes Trpm7 selectively in T cells were described by us previously (Jin et al., 2008). Mice were housed and bred in accordance with policies and protocols of the Institutional Animal Care and Use Committee (ACUC) of University of Virginia.

### Genotyping of mice

Tail samples are dissolved at 85°C for 30 minutes in 75 μl digestion buffer (25 mM NaOH, 0.2 mM EDTA), the reaction is stopped by adding 75 μl neutralization buffer (40mM Tris-HCl). One μl was used as template for PCR using MyTaq Hot Start Polymerase (Bioline; #BIO-21112) with 5x MyTaq Red Reaction Buffer (Bioline; #BIO-37112). PCR products were separated in a 1% agarose gel and visualized with ethidium bromide. The presence of *Cre recombinase* was determined using

Forward primer Cre S2F (5’-GATTTCGACCAGGTTCGTTC-3’)

Reverse primer Cre S5R (5’-GCTAACCAGCGTTTCGTTC-3’).

Presence (fl) or absence (wt) of LoxP sites flanking Exon 17 were detected using these primers:

Forward primer Geno 2F* (5’-CAGAGGTACTGGCAATTGTG-3’)

Reverse primer Geno 2R* (ACGAGGACTCAGCATATAGC-3’).

### Reagents

<u>Antibodies</u>: anti-mouse Neuropilin-1-PE (R&D Systems, #FAB5994P), rat anti-mouse CD25 PE-Cy7 (BD Pharmingen, #552880), anti-mouse CD39 PE-Cyanine7 (eBioscience, Inc. #25-0391), anti-mouse CD73 PE (eBioscience, Inc. #12-0731), rat anti-mouse CD4 FITC (BD Biosciences #553046), purified hamster anti-mouse CD28 (BD Biosciences #553294), anti-mouse/rat Foxp3 APC clone FJK-16s (eBioscience), anti-Foxp3 (eBioscience, #17-5773), purified hamster anti-mouse CD3ε chain, clone 145-2C11 (BD Biosciences #553058), anti-mouse CTLA-4 (CD152) PE (eBioscience, #12-1522), anti-mouse CD16/CD3ε antibody (Biolegend,TruStain fcX, #101320),

<u>Magnetic beads for T-cell isolation</u>: Dynabeads Untouched Mouse T Cells (Life Technologies, #11413D), CD4^+^CD25^+^Regulatory T Cell Isolation Kit mouse (Miltenyi Biotech, #130-091-041), Dynabeads FlowComp™ Mouse CD4^+^CD25^+^ T_reg_ Cells (Life technologies, #11463D)

<u>qPCR reagents and kits</u>: SensiMix SYBR Kit (Bioline USA Inc, #QT605-20), GoScript™ Reverse Transcription System (Promega Corporation, #A5001), ISOLATE II RNA Mini Kit (Bioline, #BIO-52073)

<u>Purified cytokines</u>: Human TGF-β1 (Cell Signaling Technology, #8915), human IL-2 recombinant (BD Biosciences, #BDB554603).

<u>Other chemicals</u>: CFSE (Life Tech), Concanavalin A from Canavalia ensiformis, γ-Irradiated (Sigma-Aldrich), FTY720 (Cayman Chemical), 7-AAD (BD Biosciences #559925)

### Concanavalin A induced hepatitis model

Age matched and sex matched WT and KO mice were weighed 2 hours prior to injection to control for body weight differences. Concanavalin A (40 mg/kg) or PBS was administered intravenously (i.v) through tail vein injection. Animals were monitored for 24 hours post-injection. Animals were euthanized at indicated time points or at 24 hours. Blood was collected via cardiac puncture for the measurement of serum ALT in a blinded manner. Livers were prepared as described below for subsequent analysis (blinded).

### Serum Alanine Aminotransferase (ALT) measurements

Blood was collected via cardiac puncture and centrifuged at 1500g (10min, 4°C) to collect serum. Samples with signs of hemolysis were discarded. Sera were preserved at −80°C and analyzed within 48 hours of collection. ALT activity was detected using Liquid ALT (SGPT) Reagent kit (Pointe Scientific, Inc., Canton, MI) according to the manufacturer’s instructions.

### Histology

After perfusion with PBS, a portion of liver tissue was fixed via immersion in 4% paraformaldehyde for 24 hours and transferred to 30% sucrose for 24 hours prior to tissue analysis. All tissue embedding, sectioning, and mounting were performed by the UVA Research Histology Core (University of Virginia, Charlottesville, VA). Tissues were embedded in frozen tissue blocks prior to sectioning. H&E staining was performed by the UVA Research Histology Core, and slides were photographed and analyzed on an Aperio Scanscope (Leica Biosystems, Buffalo Grove, IL).

### Immunofluorescence microscopy

Liver tissue sections were prepared on slides, rehydrated in water, washed (PBS with 0.05% Tween20), and permeabalized with PBS containing 0.1% Triton X-100 for 15 min. Slides were then blocked with blocking buffer (PBS with 1% BSA, 0.1% fish gelatin, 0.1% Triton-X-100, 0.05% Tween 20, and 5% animal serum, corresponding with the host species on the secondary antibody). Slides were stained with 1 μg/mL rat anti-mouse CD4 (Clone RM4-5, Biolegend) in blocking buffer (overnight, 4°C). Slides were then washed three times in wash buffer and stained with 0.2 μg/mL goat anti-rat AlexaFluor-594 conjugated secondary antibody (Life Technologies) solubilized in blocking buffer (2h, RT). After washing three times cells were stained with 0.2 μg/mL Hoechst 33342 nuclear dye (10 min, RT). Finally, slides were washed twice and coverslips (FischerBrand 22mm #1) were mounted over VectaMount mounting medium (Vector Laboratories, CA). Mounted samples were cured overnight at room temperature and imaged within 24 hours. All images were taken on Zeiss AxioObserver microscope with Zeiss 40X (1.3 NA) objective lens and Hamamatsu CMOS OrcaFlash 4.0 sCMOS camera. Images were then processed using NIH ImageJ software.

### Real-Time Quantitative PCR (qPCR)

Concentrations of total RNA isolated using ISOLATE II RNA Mini Kit (Bioline, #BIO-52073) were measured using NanoDrop 2000c (Thermo Scientific). cDNA was reverse transcribed using GoScript Reverse Transcription System (Promega, #A5001). qPCR experiments were always set in triplicates using SensiMix SYBR Hi-ROX Kit (Bioline, #QT605-05) in CFX connect Real-Time system (Bio-Rad, USA). After 40 cycles of PCR amplification, the data were analyzed using CFX manager 3.1 software (Bio-Rad, USA). The primers used for qPCR analysis are described below and the data were analyzed using the Δ ΔCt method (Schmittgen and Livak, 2008) using the average Ct values of βeta-2-microgobulin (β2m) to normalize for cDNA input error. Statistical analysis was based on 3 independent experiments.

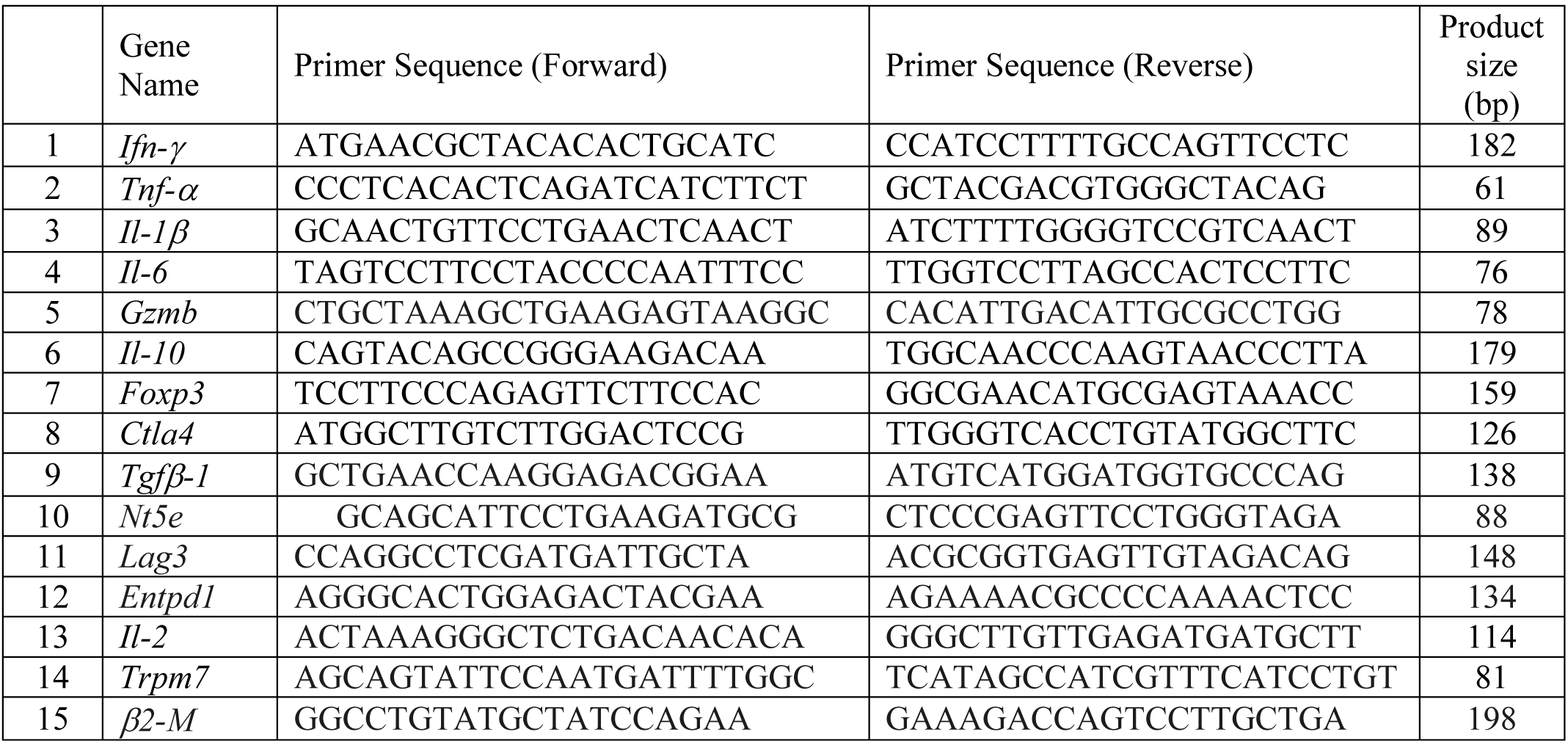

### Scanning Electron Microscopy (SEM)

Livers were perfused through the portal vein with 2.5% glutarladehyde and 4% paraformaldehyde in PBS at constant flow rate (1 mL/min). Liver sections of 5 mm thickness were removed and immersed in 2.5% glutaraldehyde and 4% paraformaldehyde (48 hours, 4°C). Samples were then washed in Osmium tetroxide (OsO_4_) and dehydrated in an ethanol series prior to critical point drying. Samples were then mounted on aluminum studs and sputter coated in gold/palladium prior to SEM on Zeiss Sigma VP HD field emission SEM microscope (Carl Zeiss AG, Oberkochen, Germany) at 3 kV.

### Isolation of intrahepatic immune cells

Livers were perfused with PBS through the portal vein. A portion of the liver was removed for immunohistological, immunofluorescent, and qPCR analysis. Livers were excised and washed with Isakov’s Modified Dulbecco’s Medium (IMDM) supplemented with 10% fetal bovine serum. Livers were then digested in 0.05% collagenase (Collagenase IV, Sigma-Aldrich, Cat. #C5138) at 37°C for 25 min. Hepatocytes were removed via centrifugation at 50xg (5 min, 4°C). Mononuclear cells (MNCs) were purified using a 40% Histodenz (Sigma-Aldrich, Cat. #D2158) gradient after centrifugation at 1500xg (20 min, 4°C). An aliquot of the total MNC was stained with trypan blue to count the live cells prior to subsequent analysis.

### Flow cytometry

Erythrocyte-free, single-cell suspensions isolated from livers, spleens or thymi were processed in FACS staining buffer composed of 150 mM phosphate-buffered saline (PBS), 0.5% Bovine Serum Albumin and 2 mM EDTA. The cells were Fc blocked with anti-mouse CD16/CD3ε antibody (Biolegend,TruStain fcX, # 101320) for 20 min and then stained with fluorochrome-conjugated antibodies. For intracellular staining of Foxp3, the cells were fixed in 1% paraformaldehyde and permeabilized with 0.1% saponin. Intracellular staining of pSTAT5 was done after concomitant fixation and permeabilization with 100% methanol. The stained cells were run on a FACSCalibur or LSRFortessa (Becton Dickinson, USA). The data was acquired by CellQuest Pro Acquisition software or FACSDiva v6 Software (Becton Dickinson, USA) respectively. Every experiment included single stain and ‘fluorescence minus one’ (FMO) controls to facilitate accurate compensation and analysis using the FlowJo version 10 software (Tree star, Ashland, OR, USA.

### Isolation of splenocytes, thymocytes and T-cell subsets

Spleens and thymi were harvested from mice euthanized by CO2 inhalation and dissected under sterile conditions. Organs were physically disrupted in cold, sterile RPMI-1640. Cell suspensions were obtained after gently passing the tissue debris through a syringe (18-ga needle). Cell suspensions were filtered through 70 μM nylon mesh and the contaminating erythrocytes were lysed using ACK buffer (150 mM NH4Cl, 10 mM KHCO_3_, 100 μM Na2EDTA; pH 7.3). These cell suspensions were then subjected to appropriate magnetic separation to isolate different subsets. We followed the protocols included in these kits: Dynabeads Untouched Mouse T Cells (Life Technologies, #11413D), CD4^+^ CD25^+^Regulatory T Cell Isolation Kit mouse (Miltenyi Biotech, #130-091-041), Dynabeads FlowComp™ Mouse CD4^+^ CD25^+^ T_reg_ Cells (Life technologies, #11463D).

### T-cell activation

Freshly isolated T-cell subsets were cultured in 96-well round bottom plates. Cell density was 0.5 ×; 10^6^/ml and culture volume was 200 μL/well. The cells were activated with Concanavalin A (5 μg/ml) for 48h. Alternatively, cells were activated using a mixture of anti-CD3ε (2.5 μg/ml) and anti-CD28 (1 μg/ml) for 48h. After 48h of activation, cells or the supernatants were harvested for analysis. For CFSE proliferation assays, the T-cell subsets were cultured at a reduced density (50000/well) in 96 well round bottom plates. Proliferation was then stimulated using a mixture of anti-CD3ε (0.25 μg/ml) and anti-CD28 (0.125 μg/ml) and analysis was carried out after 3d of culture.

### Transwell-migration assay

Each transwell (5.0 μm pore size membrane) from Corning costar incorporation contained 5×10^5^splenic T cells from WT or KO, suspended in 100 μl of complete medium (top chamber). Transwell inserts were placed in a 24-well tray containing 600 ml of variably diluted conditioned medium containing chemotactic signals secreted by LPS-treated bone marrow derived macrophages (BMDM). During the incubation of Transwell plates (16 h, 37° C, 5%CO2), the T cells transmigrated to the bottom chamber. They were manually counted using trypan blue. Data are represented as a percentage of T cells that migrated across the membrane.

### Luminex multiplex and ELISA assays

Cell culture supernants were thawed from −80°C and were run in duplicate to measure the cytokine levels. The data were acquired and analyzed using the Luminex multiplex assay (Luminex 100 IS). Standards were assayed for each cytokine with QC1 and QC2 in duplicates with beads count of <25. The cytokine concentrations were calculated by curve-fitting five concentrations of standards. ELISA assays were carried out using instructions included in the kits by the manufacturer.

### Electrophysiology

TRPM7 current (I_TRPM7_) were measured in the whole cell configuration as illustrated **(Fig S4F)**. T cells were activated with a mixture of anti-CD3ε and anti-CD28 antibody (or Concanavalin A) and cultured in 96-well plate (37°C, 5% CO2) for 1-4 days before patch-clamp electrophysiology. The standard external solution contained (in mM): 135 Na-methanesulfonate (Na-MeSO_3_), 5 Cs-gluconate, 2.5 CaCl_2_, 10 HEPES, pH 7.3 (adjusted with NaOH), Osmolality 280-290 mOsm/Kg. The standard pipette solution contained (mM) 110 Cs-gluconate, 0.5 NaCl, 0.75 CaCl_2_, 10 HEPES, 10 HEDTA, 1.8 Cs_4_-BAPTA, 2 Na_2_ATP, pH 7.3 (adjusted with CsOH). Osmolality was 273 mOsm/Kg and calculated free [Ca^2+^] = ∼100 nM. We used the maxchelator algorithm to calculate free Ca^2+^: http://maxchelator.stanford.edu/webmaxc/webmaxcS.htm FTY-720 (2 μM) or MgCl_2_ (10 mM) was used with external solution to inhibit TRPM7 currents. All currents were recorded using an Axopatch 200B amplifier (Molecular devices, Sunnyvale, CA). The recording protocol consisted of 400 ms ramps from −100 mV to +100 mV and holding potential (HP) of 0 mV. The signals were low-pass filtered at 5 kHz and sampled at 10 kHz. All electrophysiology experiments were done at RT (∼22°C). The average current densities were plotted with the relevant statistical information as a box chart (see *Statistics* below).

### T-cell proliferation suppression assay

Splenic cell suspensions were isolated from WT and KO mice. Effector T cells (T_eff_) or regulatory T cells (T_reg_) were then isolated through negative and positive magnetic selection using Dynabeads Flow Comp Mouse CD4+ CD25+ T_reg_ Cells kit (Lifetech). T_eff_ cells were labeled with 5 μM CellTrace CFSE (Carboxyfluorescein succinimidyl ester) for 20 min, washed and cultured in RPMI-1640 medium supplemented with 2mM L-glutamine, 10% FBS, 50 μM β-mercaptoethanol, 1% sodium pyruvate, 5mM HEPES, 100 U/ml penicillin and 100 μg/ml streptomycin. The CFSE-labeled T_eff_cells were cultured in 96-well round bottom plate and stimulated with a mixture of anti-CD3ε and anti-CD28 antibodies in 5% CO_2_ humidified incubator (37°C, 3 days). The number of T_eff_ cells was kept constant at 5 ×; 10^4^ cell per well, whereas the number of cocultured T_reg_ cells varied for indicated ratios. On the 4^th^ day the cells were collected and stained for 7-AAD, CD4 and CD25 before flow cytometry analysis on FACSCalibur instrument. The data were analyzed using FlowJo software and represented as the percentage of T_eff_ cells showing proliferation-induced dilution of CellTrace CFSE dye. This parameter reflects the percentage of cells undergoing cell division.

### *Ex vivo* generation of regulatory T cells from thymocytes

Thymocytes were isolated from WT and KO as single cell suspensions and cultured in X-vivo 15 growth medium (Sartorius stedim Biotech, Cat #04-744Q) supplemented with 100 U/ml penicillin, and 100 μg/ml streptomycin. Thymocytes were activated with anti-CD3ε antibody in presence of recombinant hIL-2 (10 ng/ml) in 96-well round bottom plates (5% CO_2_, 37°C, 48 h). After 48h, thymocytes were collected and were either used to isolate mRNA for Foxp3 mRNA analysis by qPCR or analyzed by flow cytometry for immunophenotyping and intracellular Foxp3 expression.

### Chromatin Immunoprecipitation (ChIP) - qPCR

An equal number of freshly isolated thymocytes from WT and KO mice were cross-linked (1% formaldehyde, 10 min) before stopping the cross-linking reaction with glycine. The cytoplasmic contents were extracted and removed using cell lysis buffer containing 1% SDS. The nuclear pellets were resuspended in ChIP buffer (1.1% TritonX-100 containing nuclear lysis buffer). These nuclear pellets were subjected to sonication using Bioruptor Pico (Diagenode Inc., NJ, USA). The sheared chromatin was then immunoprecipitated (12h, 4° C) using ChIP grade antibodies (anti-pSTAT5 (Tyr694) and control-IgG) and ChIP grade protein G magnetic beads (Cell signaling Technologies). The immunoprecipitated ChIPs were then eluted with ChIP elution buffer and treated with proteinase K (12h. 65°C) to release the DNA fragments. The released DNA was purified using column based DNA purification kit (Qiagen) and subjected to qPCR using the primers shown below. The data were then analyzed to assess the occupancy of STAT5 on IL2Ra promoters in KO thymocytes, relative to WT thymocytes. Briefly, the Ct values obtained using mock IgG ChIPs were first used to calculate the fold change in anti-STAT5 ChiPs in WT and KO samples. The KO STAT5 ChIPs showed significantly higher fold change values compared to WT – these were shown as bar graphs after normalizing the WT values to 1.

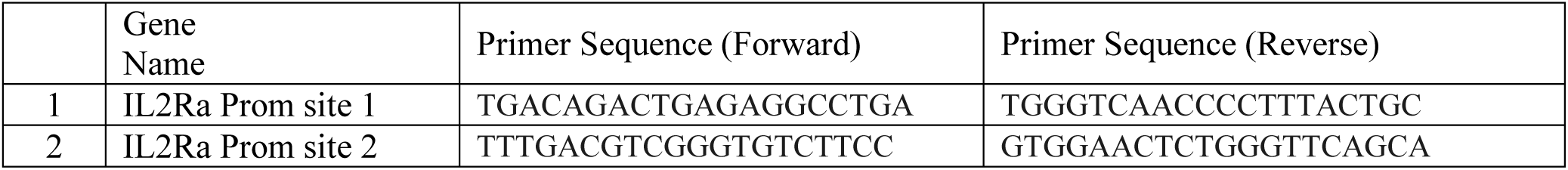

## LEGENDS for Supplementary figures

**No Figure S1** *because all supplementary figures are related to their cognate figures and Fig. 1 does not point to any supplementary data*

**Figure S2.**
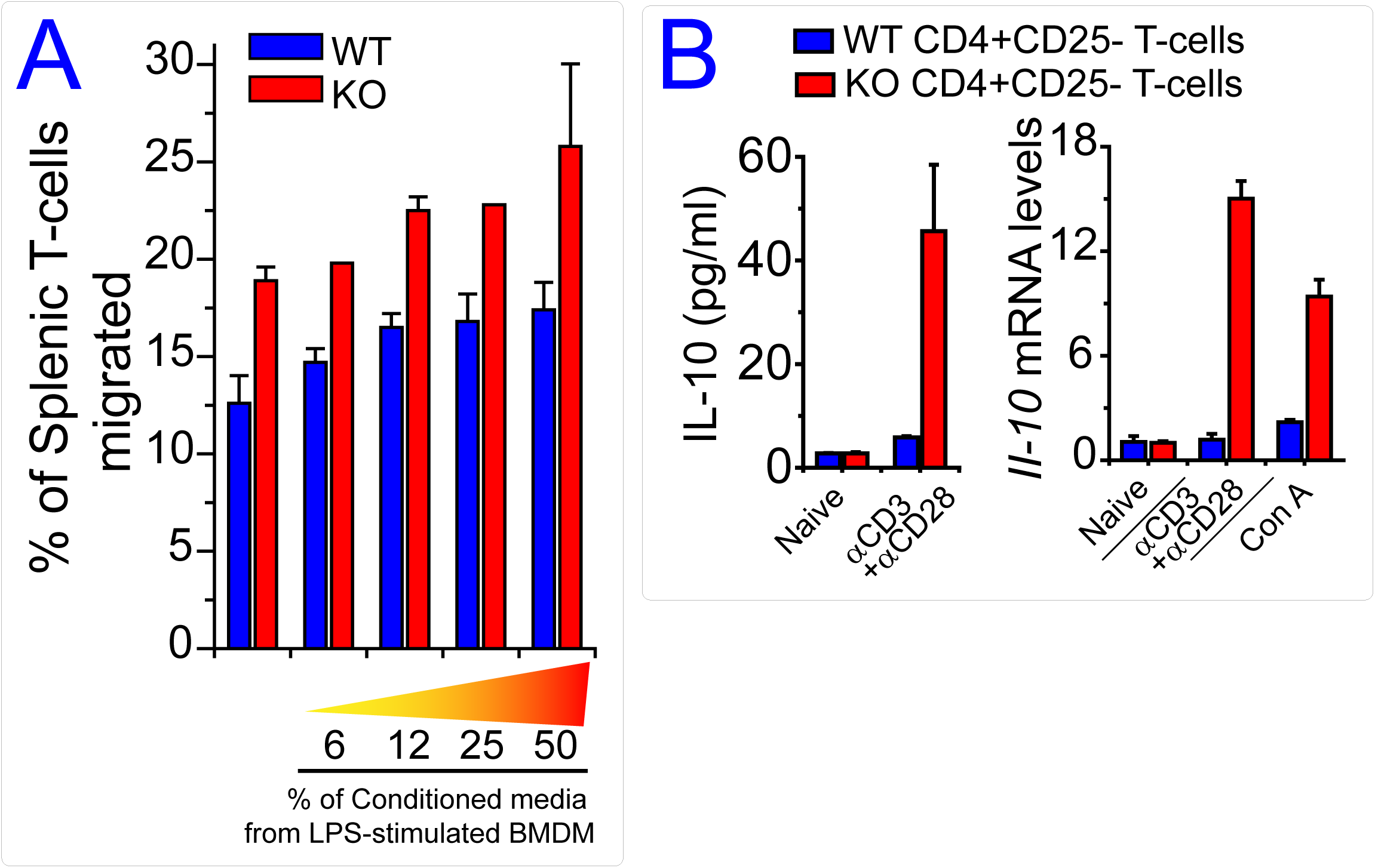
Related to Figure 2. **(A)** Transwell migration of WT and KO T cells through membrane (pore size: 5.0 μm). Conditioned medium from LPS-treated bone marrow derived macrophages (BMDM) was used as chemoattractant in the bottom chamber (see methods for details). Data are represented as a percentage of T cells that migrated across the membrane. **(B)** *Left panel* shows quantification of IL-10 secretion by activated WT and KO CD4+ T cells (anti-CD3 and anti-CD28 antibodies, 72h). Bar graphs represent mean cytokine levels (n=3); error bars reflect SEM. *Right panel* shows the quantification of IL-10 mRNA in WT and KO CD4+ T cells after 48h of activation.

**Figure S3.**
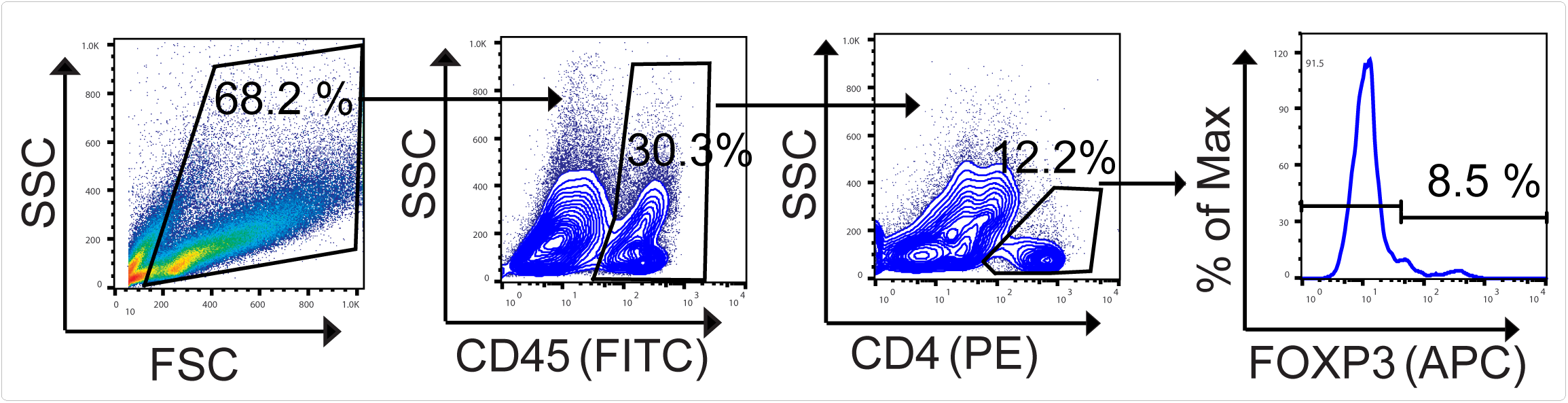
Related to Figure 3. Gating scheme used to derive the Foxp3 histograms shown in *figure 3F*.

**Figure S4.**
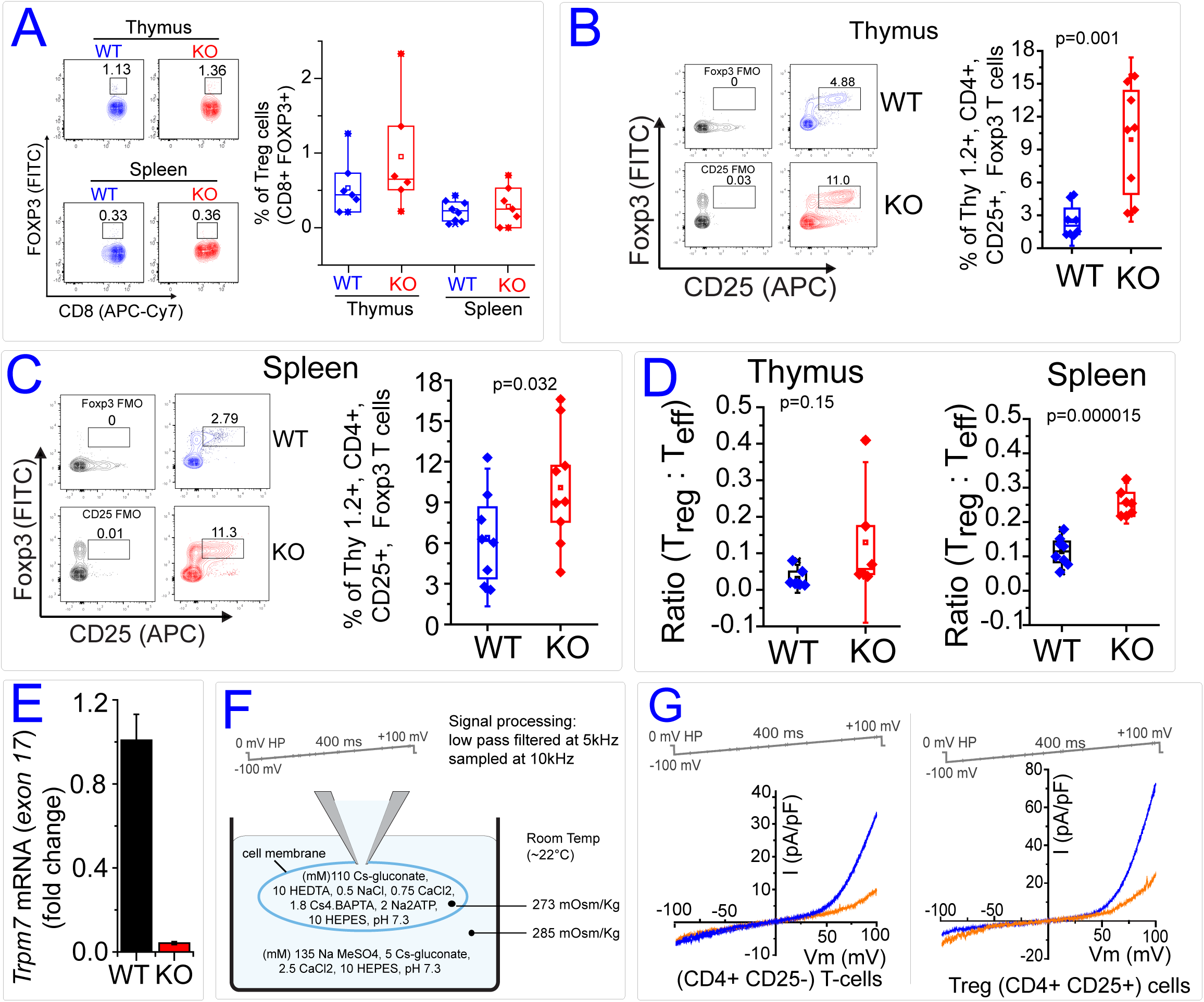
Related to Figure 4. **(A)** Bivariant cytograph *(Left)* shows the percentage of Foxp3+CD8+ cells in the thymi and spleens of WT and KO mice. Box chart *(Right)* shows the statistics and quantification of CD8+Foxp3+ T cells in thymi and spleen. **(B)**. Flow cytometry contour plot *(Left)* shows the staining of cell surface CD25 and intracellular FOXP3 in previously gated Thy1. CD4+ WT and KO thymocytes. The box chart *(Right)* quantifies the percentage of CD4+CD25+Foxp3+ thymocytes in WT and KO mice. The mean value (empty square) and the median (horizontal line) are shown in the boxes. The data were collected from four independent experiments and the diamonds in the box chart denote individual mice. The indicated p value was calculated using a two-tailed student’s t test. **(C)**. Flow cytometry contour plot *(Left)* shows the staining of cell surface CD25 and intracellular FOXP3 in previously gated Thy1.2+ CD4+ splenic T cells obtained from WT and KO mice. The box chart *(Right)* quantifies the percentage of CD4+CD25+Foxp3+ T cells in the WT and KO spleens. The mean value is shown by an empty square and the median by a horizontal line. The data were collected from four independent experiments and the diamonds in the box chart denote individual mice. The indicated p value was calculated using a two-tailed student’s t test. **(D)**. The box chart shows the calculated ratios of T_reg_: T_eff_ cells in the thymi (Right) and spleens of WT and KO mice *(Left)*. The indicated p value was calculated using a two-tailed unpaired student t-test. **(E)**. Bar graph shows the expression of *Trpm7* T_reg_ cells (CD4+CD25+) obtained from WT and KO spleens. The qPCR primers were directed at the exon 17 of *Trpm7* and the the relative reduction of TRPM7 exon 17 containing mRNA in the KO T_reg_ cells is shown. **(F)**. Whole cell patch clamp recording conditions used to isolate and measure ITRPM7 are schematized. Internal (pipet) and external (bath) solutions are shown. Voltage ramp parameters used to derive the I-V relationship of I_TRPM7_ are shown along with filtering and sampling parameters used for signal processing. **(G)**.IV relationship (blue trace) of I_TRPM7_ in WT CD4+CD25-(T_conv_) and CD4+CD25+ (T_reg_) cells obtained by whole cell patch clamp recordings. Orange trace shows the inhibition of ITRPM7 by perfusion of 10 mM Mg^2+^.

**Figure S5.**
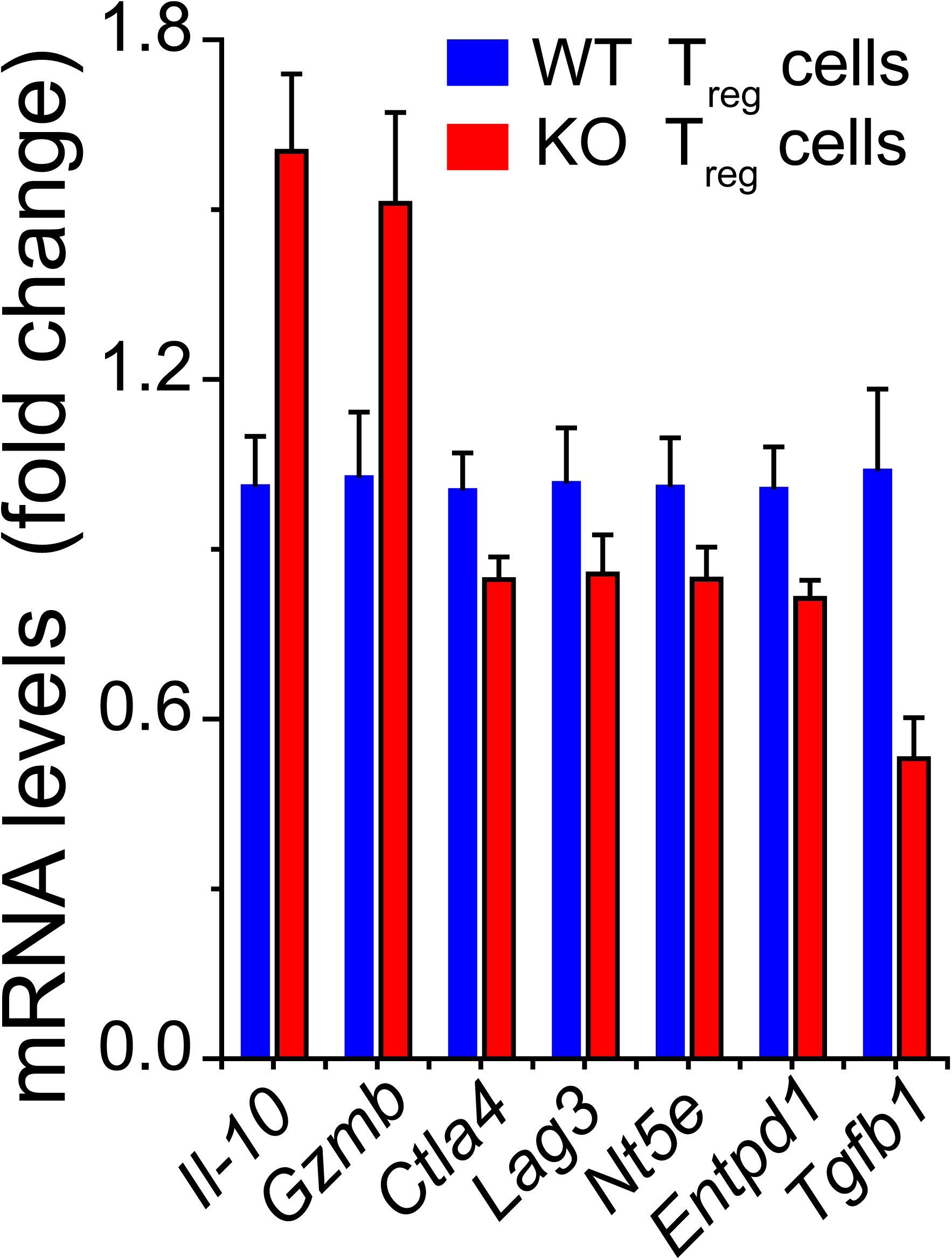
Related to Figure 5. Gene expression (qPCR) of T_reg_ markers in freshly isolated WT (n=4) and KO (n=6) splenic CD4+CD25+ cells.

**Figure S6.**
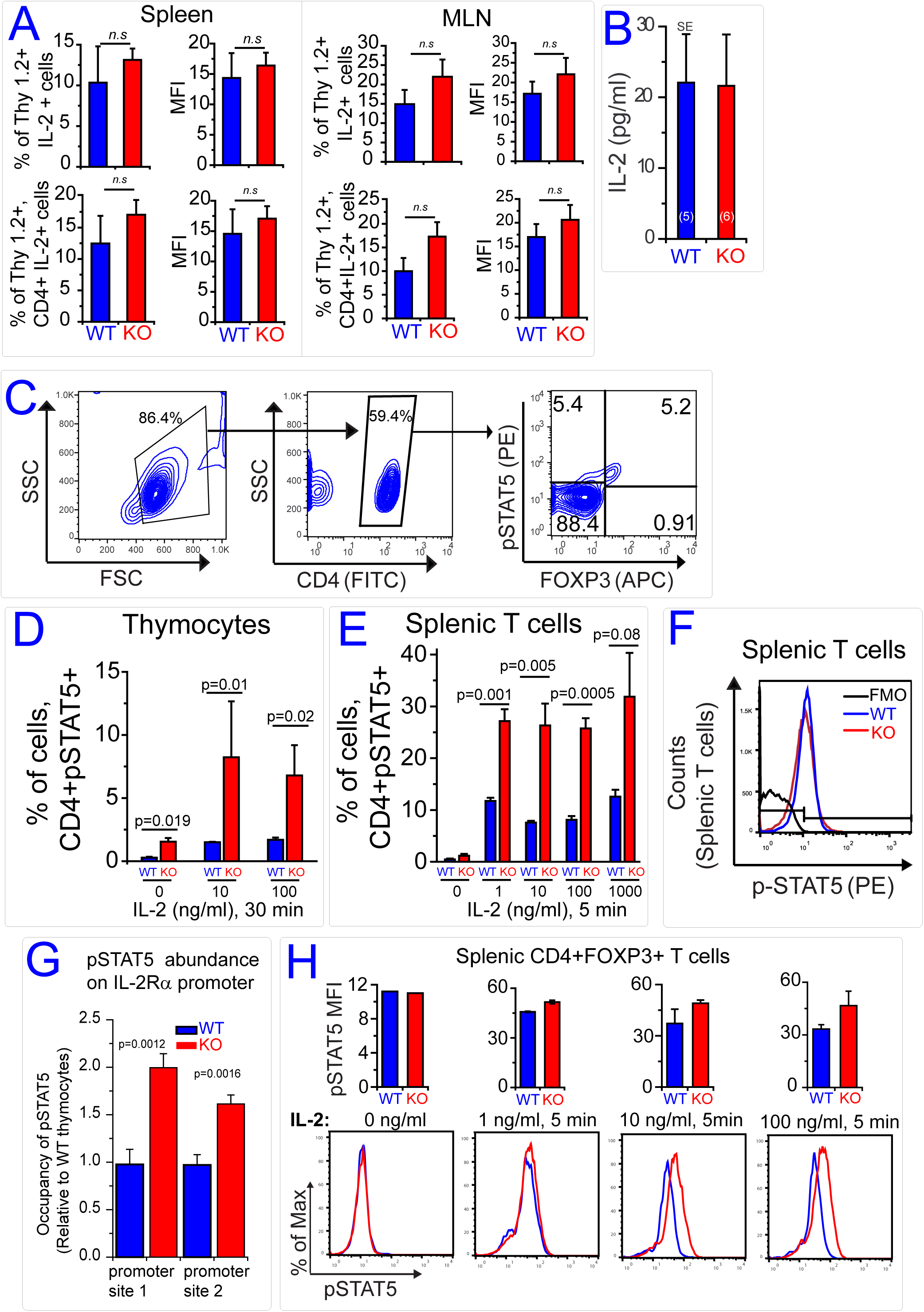
Related to Figure 6. **(A)**. The bar graphs show the expression on IL2 in T cells obtained from WT and KO spleens *(Left)* and mesenteric lymph nodes *(Right).* In each case bar graphs in the *upper panels* show the analysis of total T-cells (Thy1.2+) and the bar graphs in the *lower panels* show the analysis of CD4+ T cells (Thy1.2+ CD4+). Both the percentage of IL2-expressing cells and MFI of IL2 staining are quantified. **(B)**. The bar graph shows the average IL2 concentrations, acquired using ELISA, in sera obtained from WT (5 mice) and KO (6 mice). The data graphs reflect SEM. **(C)**. Representative gating strategy used to analyze CD4+ cells for pSTAT5 and Foxp3 expression is shown. **(D)**. Bar graph shows the percentages of pSTAT5+ thymocytes as determined by the addition of upper left and upper right quadrant gates from flow cytometry analysis shown in *Figure 6I*. The error bars reflect SEM (n=4); p values were calculated using t-test. **(E)**. Bar graph shows the percentages of pSTAT5+ splenic T cells as determined by addition of upper left and upper right quadrant gates from flow cytomtery analysis shown in *Figure 6K*. Error bars reflect SEM (n=4); P-values calculated using t-test. **(F)**. Flow cytometry histogram shows intracellular staining of phospho-STAT5 (pSTAT5) in splenic T cells isolated from WT and KO mice. Fluorescence minus one (FMO) control reflects background staining in WT lymphocytes without anti-pSTAT5 antibody. **(G)**. Bar graph shows ChIP-qPCR analysis of pSTAT5 occupancy on two different regions of IL2Rα promoter in WT and KO thymocytes. The occupancy seen in WT thymocytes was normalized (blue bars) and relative difference in KO occupancy is shown (red bars). **(H)**. The bar graphs compare the MFI of pSTAT5 in splenic CD4+FOXP3+ T cells obtained from WT and KO mice. Representative overlaid histograms of pSTAT5 staining in corresponding lower panels.

**Figure S7.**
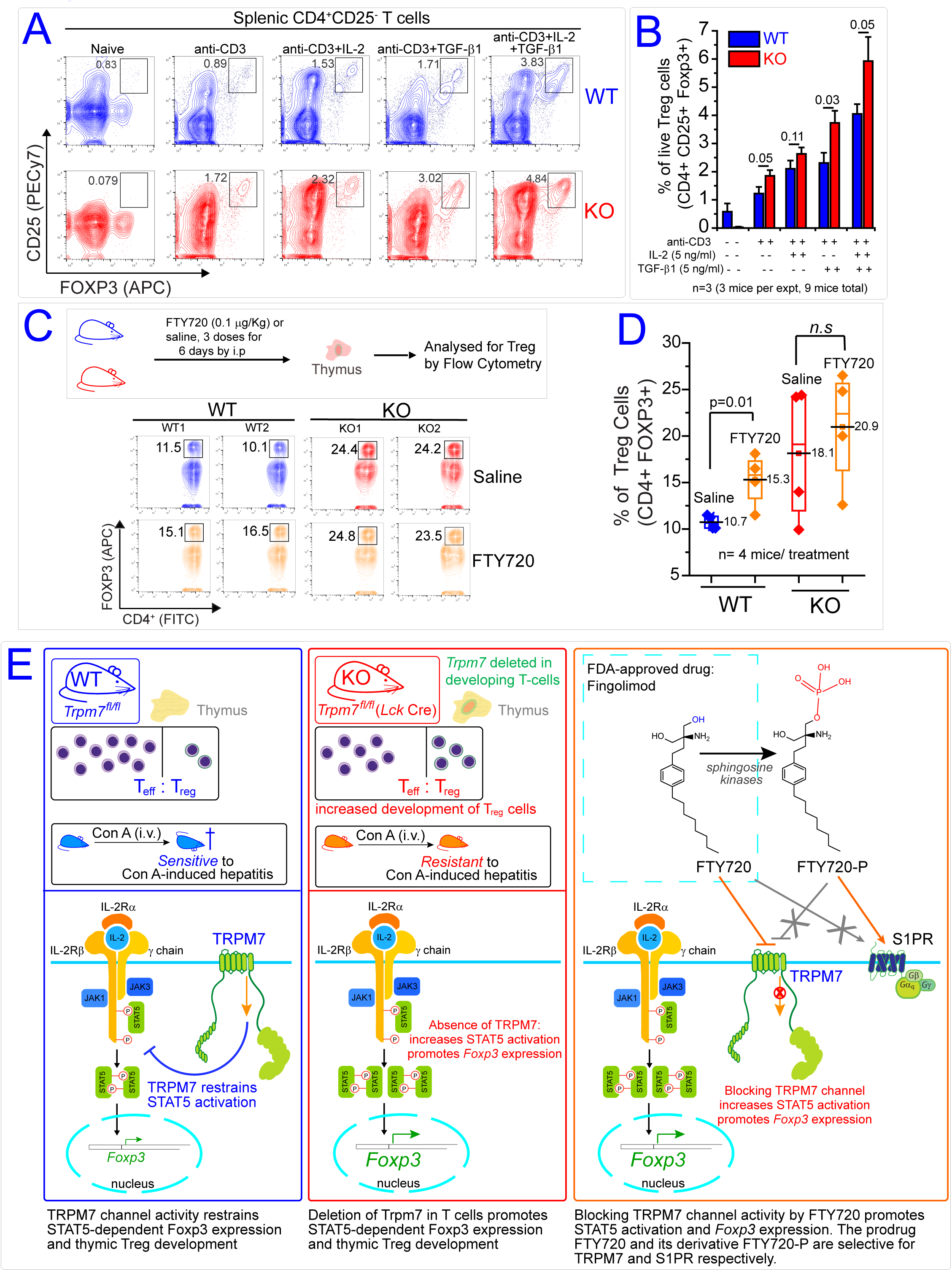
Related to Figure 7. **(A)**. Bivariant contour plots show the induction of CD25 and FOXP3 in WT and KO splenic T cells (CD4+CD25-) in response to indicated activation treatments. The cells were initially gated for viability. The data are representative of 3 independent experiments. **(B)**. Bar graphs show the mean percentage of T_reg_ cells (CD4+CD25+FOXP3+) induced in the indicated culture conditions. Error bars reflect SEM (n=3). Each independent experiment used T cells pooled from 3 mice. **(C)**. Upper panel shows the experimental scheme. WT (blue) and KO (red) mice received 3 doses of FTY720 (0.1μg/Kg body weight; i.p.) for 6 days by intraperitoneal injection. The bivariant contour plots (lower panels) show typical results of thymocytes analyzed to quantify CD4+FOXP3+ T_reg_ cells. **(D)**. Box charts show the mean (horizontal line) and scatter of the percentages of CD4+FOXP3+ T_reg_ cells found in the thymi of saline and FTY720-treated WT and KO mice. The p values were calculated using two-tailed unpaired student t-test and filled diamonds represents individual mice. **(E)**. Illustrated summary of overall findings and proposed model.

